# Duplicate gene retention is partly repeatable across independent whole-genome duplications in plants, fungi and animals

**DOI:** 10.64898/2026.08.29.748011

**Authors:** Krishnendu Sinha, Nabanita Ghosh

## Abstract

Whole-genome duplication (WGD) has recurred throughout eukaryotic evolution. After each event most duplicated gene pairs (ohnologs) have returned to single copy, but a non-random subset has been retained. Whether the same ancestral genes are retained again after independent WGDs, and why, has not been quantified gene by gene. Using hierarchical orthologous groups, we follow ancestral genes through seven independent WGDs in plants, fungi and animals (42 descendant genomes). A gene retained after one reference angiosperm WGD is 1.67–1.73 times as likely to be retained after another. A liability-threshold model attributes 29% (95% CI 25–36%) of gene-level variation in retention propensity to a component shared across WGDs, and 76 gene features account for 33–40% of the association between WGDs. In prespecified tests with predictions hash-recorded before outcomes were examined, a score from three angiosperm WGDs predicts retention after the apple–pear and budding-yeast WGDs (Spearman *ρ* = 0.169 and 0.280; *P <* 0.001), whereas the teleost test does not meet its criterion. Comparing each gene across genomes descended from one WGD, holding the gene fixed, shows that yeast genomes lose physical complex partners together (*z* = 8.1), as dosage balance predicts, whereas plant and fish genomes lose neighbouring genes together, a lineage-specific process that does not recur between events. Sequence constraint measured before duplication accounts for 30% of the plant-to-yeast transfer. Thus a gene-intrinsic propensity linked to dosage balance and constraint recurs across kingdoms, whereas deletions and lineage history decide where losses fall.

## Introduction

Whole-genome duplication (WGD), or polyploidy, doubles every gene of a genome in a single step. It has recurred throughout eukaryotic history: all flowering plants descend from several ancient WGDs (Jiao et al. 2011; Van de Peer et al. 2017), as do the baker’s yeast lineage (Wolfe and Shields 1997; Kellis et al. 2004) and teleost fishes (Jaillon et al. 2004), and many crops, including wheat, cotton, potato and strawberry, are recent polyploids. The paired gene copies produced by WGD, called ohnologs, are a major source of evolutionary novelty (Ohno 1970; Conant and Wolfe 2008). Yet the duplicated state is usually transient. Over the following millions of years one copy is lost from most ohnolog pairs, a process called fractionation, and only a minority of pairs are retained (Van de Peer et al. 2017; Scannell et al. 2006).

Retained ohnologs are not a random sample. In plants, yeasts and vertebrates they are enriched for transcription factors, signalling proteins, ribosomal proteins and subunits of protein complexes (Maere et al. 2005; Blanc and Wolfe 2004; Seoighe and Gehring 2004; Papp et al. 2003; Makino and McLysaght 2010). The dosage-balance hypothesis explains this bias: proteins that act in fixed stoichiometry are doubled together by WGD, and loss of a single copy would unbalance their complexes or pathways (Birchler and Veitia 2007; Freeling 2009; Edger and Pires 2009). Other ohnologs persist because the copies partition or diversify ancestral functions (Force et al. 1999; Lynch and Conery 2000). Fractionation is further shaped by the event itself, for example by which parental subgenome comes to dominate after allopolyploidy (Thomas et al. 2006; Schnable et al. 2011; Cheng et al. 2018). Gene-family studies have noted that families retained after one WGD are often retained after another (Paterson et al. 2006; Barker et al. 2008; Li et al. 2016; Mandáková et al. 2017), and that some families almost always return to single copy (De Smet et al. 2013).

These observations bear on a general question about the repeatability and predictability of evolution (Gould 1989; Blount et al. 2018). Independent WGDs in unrelated lineages are natural replicates of one evolutionary experiment, because each duplicates, among others, the descendants of the same ancient ancestral genes. Whether the retention of a particular ancestral gene after one WGD carries information about its retention after another has, however, been addressed mainly through categories of genes. Three questions therefore remain open: how large the repeated component is at the level of individual ancestral genes; whether it is explained by gene properties such as function or dosage sensitivity; and whether it predicts retention in WGDs that were not used to build the prediction. Answering these questions requires identifying the same ancestral gene across events separated by up to 1.5 billion years, treating comparisons that share genes and events as dependent, and allowing for the unequal numbers of genomes that represent each event.

Here we follow the same ancestral genes through seven independent WGDs, in the mustard, grass, legume and apple–pear lineages, in budding yeasts, in teleost fishes and in stylommatophoran land snails, using published reconstructions that record ohnolog presence in every descendant genome (Hao et al. 2022; Conant 2023) and hierarchical orthologous groups to identify ancestral genes (Altenhoff et al. 2013, 2024). We find that retention after one WGD is consistently associated with retention after another; that about 30% of gene-level variation in retention propensity is shared across WGDs; that 76 gene properties account for about one-third of this shared component; and that predictions from angiosperm WGDs, fixed and hash-recorded before the target data were examined, succeed in the apple–pear and budding-yeast WGDs. We then compare each gene across the genomes descended from one WGD, a design that holds the gene itself fixed, and find that yeast genomes lose protein-complex partners together, whereas plant and fish genomes lose neighbouring genes together (Table 1). Technical terms are defined in a glossary (Supplemental Note S9).

**Table 1.** Main findings. CI, 95% confidence interval from bootstrap resampling of HOGs; OR, odds ratio.

| Question and measure | Estimate | HOGs |
| --- | --- | --- |
| *Is retention after one WGD associated with retention after another?* |  |  |
| Retained after legume WGD, if retained / lost after At-α | 50% / 29% (RR 1.73, CI 1.52–1.98) | 1,445 |
| OR, reference WGD pairs | 2.17–2.45 (all CIs exclude 1) | 636–1,445 |
| Retained after all three reference WGDs | 28 observed vs 16.2 expected ($P = 0.0008$) | 337 |
| *How much of retention propensity is shared across WGDs?* |  |  |
| Repeatability (liability-threshold model) | 29% (CI 25–36%) | 20 genomes |
| Observed correlation as share of attainable maximum | 24–32% | 3 pairs |
| *Is it explained by gene properties?* |  |  |
| Share of association explained by 76 features | 33% (cross-fitted) to 40% (in-sample, chance-corrected) | 3 pairs |
| Adjusted OR, reference WGD pairs | 1.85–2.09 | 3 pairs |
| *Does it predict independent WGDs? (prespecified)* |  |  |
| Apple–pear: Spearman $\rho$; top vs bottom quartile | 0.169 ($P = 0.00047$); 35% vs 21% | 373 |
| Budding yeast: Spearman $\rho$; top vs bottom quartile | 0.280 ($P = 0.00009$); 43% vs 17% | 186 |
| TGD: Spearman $\rho$ | 0.107 ($P = 0.094$); criterion not met | 151 |
| *Does it extend across kingdoms?* |  |  |
| TGD and yeast WGDs (no plant data): OR | 2.10 (CI 1.29–3.37) | 506 |
| Plant and yeast WGDs: latent correlation | 0.32–0.41 | 234–449 |
| *What acts within genomes? (exploratory)* |  |  |
| Yeast: coupling of a gene’s loss to its complex partners’ loss ($\theta$) | 0.31 (CI 0.22–0.41; $z = 8.1$) | 697 |
| Plants and fishes: same coupling ($\theta$, upper 95% bound) | $\leq 0.10$ | 5 WGDs |
| Plants and fishes: coupling to the adjacent gene ($\theta$) | 0.08–0.32 ($z \geq 7.5$) | 5 WGDs |
| Plant-to-yeast transfer explained by pre-WGD constraint | 30% | 630 |

## Results

### Seven independent WGDs and a common set of ancestral genes

We analysed seven WGDs that occurred on separate branches of the eukaryotic tree between about 38 and 277 million years ago (Fig. 1A; Supplemental Tables S9–S10). Three angiosperm WGDs, the Brassicaceae At-*α* event (six descendant genomes), the Poaceae grass *ρ* event (five) and the papilionoid legume event (four), formed the reference set from which predictions were built. The Maleae (apple–pear) WGD (five genomes), the budding-yeast WGD (eleven) and the teleost genome duplication (TGD; eight) served as

**Figure 1.**
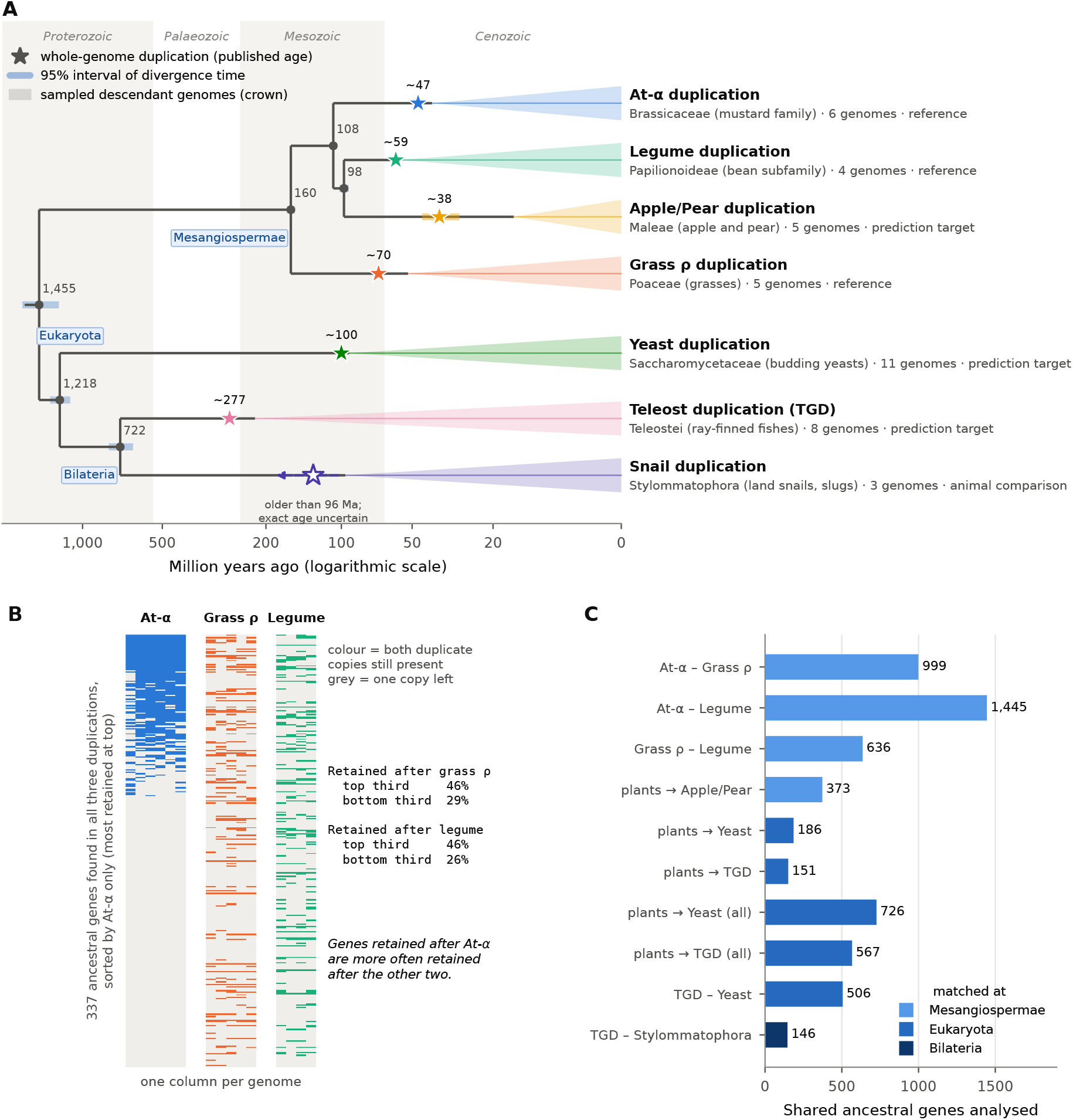
Seven independent WGDs and the ancestral genes followed through them. **A** Dated tree of the lineages studied. Stars mark WGDs at their published ages (bars, published ranges); grey numbers are divergence times in millions of years (TimeTree 5 medians, with 95% intervals as blue bars); coloured wedges span the crown age of the sampled descendant genomes. The stylommatophoran WGD predates the sampled crown but its exact age is uncertain (open star). Blue labels mark the ancestors at which HOGs were defined for matching genes across WGDs. Time axis, log_10_(*t* + 10). **B** Raw retention data for the 337 HOGs present in all three reference WGDs. Rows are HOGs and columns descendant genomes; a coloured cell indicates that both ohnologs are present. Rows are sorted only by retention occupancy in the At-*α* WGD. **C** Numbers of HOGs shared in each comparison and the taxonomic level at which they were matched; “all” denotes the enlarged populations of the sensitivity analysis.

prediction targets, and a WGD shared by stylommatophoran land snails and slugs (three genomes), reconstructed in this study, provided an animal comparison independent of plant data.

For each WGD, the POInT reconstructions represent every ancestral locus as a “pillar” of two duplicate tracks followed through all descendant genomes, so that each descendant either retains both ohnologs or has lost one (Hao et al. 2022; Conant 2023). We summarized each pillar by its retention occupancy, the fraction of descendant genomes that retain both ohnologs, and by a binary outcome. A pillar was classed as retained after a WGD when at least one descendant retained both ohnologs; 27–42% of pillars met this criterion in the At-*α*, grass *ρ*, legume, yeast and TGD events (Supplemental Table S10). In the young Maleae event 97% of pillars retain both ohnologs in at least one of five closely related genomes, so the informative contrast there is complete retention in all five genomes (27% of pillars). Rank-based analyses use the full occupancy distribution and do not depend on either threshold.

To recognize the same ancestral gene in different WGDs we used hierarchical orthologous groups (HOGs) from OMA (release All.May2026) (Altenhoff et al. 2013, 2024). A HOG collects all genes descended from a single gene of a named ancestor; we used the ancestors of mesangiosperms for comparisons among plants, of eukaryotes for comparisons involving yeast and the TGD, and of bilaterian animals for the TGD– snail comparison (blue labels in Fig. 1A). A pillar was assigned to a HOG only when proteins from at least two descendant species matched OMA proteins exactly, all matches supported the same HOG, and the pillar–HOG relationship was one-to-one within the event (Methods). These rules assigned 69–79% of reference pillars, giving 8,631 mesangiosperm HOGs, of which 636 to 1,445 were shared by each pair of reference WGDs and 337 by all three (Fig. 1C). The deep comparisons involve fewer genes (107–506 per pair), chiefly because a mesangiosperm HOG can be lifted one-to-one to a eukaryotic HOG only when no other angiosperm lineage descends from the same eukaryotic gene; the effect of these exclusions is examined below.

The central pattern is visible in the raw data (Fig. 1B). When the 337 HOGs present in all three reference WGDs are sorted solely by their retention after At-*α*, the top third were retained after the grass *ρ* WGD in 46% of cases and the bottom third in 29%; the corresponding values for the legume WGD are 46% and 26%.

### Retention after one WGD predicts retention after another

Retention was positively associated in every comparison among the reference WGDs (Fig. 2A; Supplemental Table S1). Of the HOGs retained after At-*α*, 50% were also retained after the legume WGD, compared with 29% of those that returned to single copy after At-*α* (risk ratio [RR] 1.73, 95% CI 1.52–1.98; odds ratio [OR] 2.45, 1.97–3.07; one-sided Fisher’s exact *P* = 9.5 *×* 10*^−^*^16^; *n* = 1,445). The corresponding values were 43% versus 26% for At-*α* and grass *ρ* (RR 1.67, 1.40–2.01; OR 2.17, 1.66–2.88; *P* = 1.9 *×* 10*^−^*^8^; *n* = 999) and 49% versus 28% for grass *ρ* and legume (RR 1.73, 1.41–2.14; OR 2.42, 1.72–3.46; *P* = 3.9 *×* 10*^−^*^7^; *n* = 636). Retention after one reference WGD discriminated retained from lost HOGs in another with an area under the ROC curve (AUC) of 0.60–0.62.

**Figure 2.**
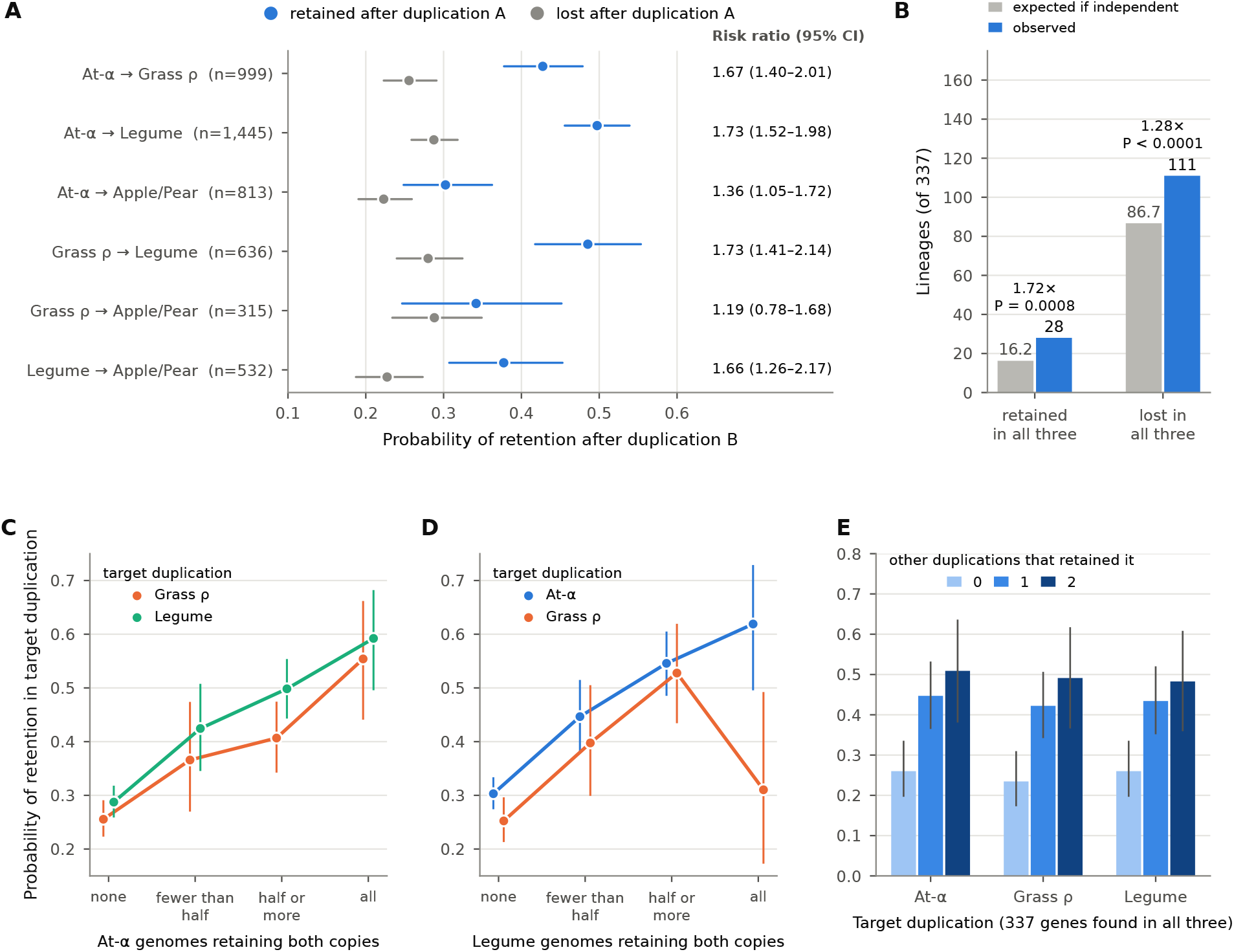
Retention after one WGD is associated with retention after another. **A** Probability of retention after WGD B for HOGs retained (blue) or returned to single copy (grey) after WGD A, with 95% bootstrap intervals; right, risk ratios. Retention: both ohnologs present in at least one descendant genome (apple–pear: in all five). **B** HOGs retained, or returned to single copy, after all three reference WGDs compared with the number expected under independence; *P* , permutation test. **C, D** Probability of retention in a target WGD by the number of At-*α* (C) or legume (D) genomes retaining both ohnologs. **E** Probability of retention in each reference WGD by the number of the other two reference WGDs that retained the HOG.

Associations with the apple–pear WGD, whose outcome is complete retention in five genomes, were in the same direction. Complete retention was more frequent among HOGs retained after the legume WGD (38% versus 23%; OR 2.06, 1.38–3.07; *P* = 0.0003; *n* = 532) and after At-*α* (30% versus 22%; OR 1.51, 1.07–2.11; *P* = 0.011; *n* = 813). The association with grass *ρ* was weaker and not statistically supported in the smallest shared set (34% versus 29%; OR 1.28, 0.71–2.19; *P* = 0.223; *n* = 315).

On the rank scale, which uses the full occupancy distribution, the correlations of event-wide retention percentiles between reference WGDs were 0.188 (At-*α*–grass *ρ*), 0.233 (At-*α*–legume) and 0.193 (grass *ρ*– legume) in the prespecified analysis (recomputation from the released tables differs only in the third decimal place; Supplemental Table S1). Their sample-size-weighted Fisher mean, *T*_replay_ = 0.210 (95% CI 0.172– 0.248; permutation *P* = 1/100,001), was the first prespecified test of the study and met its decision criteria (Table 2).

**Table 2.** Prespecified tests, decision rules and outcomes. Decision rules were fixed and the prediction or analysis population hash-recorded before the association was computed (Methods; freeze records in Supplemental Table S11). Statistics are one-sided permutation tests; intervals are 95% bootstrap intervals. *T* , (*n −* 3)-weighted Fisher mean of pairwise correlations; *ρ*, Spearman correlation; *r*, Pearson correlation of event-wide percentiles. Reference: At-*α*, grass *ρ* and legume WGDs.

| Test | HOGs | Decision rule (fixed in advance) | Result | Decision |
| --- | --- | --- | --- | --- |
| Retention association among reference WGDs | 2,406 | $T > 0$, $P \leq 0.001$, lower bound $> 0$; $T \geq 0.10$ | $T = 0.210$ (0.172–0.248),<br>$P = 1/100,001$ | Met |
| Each reference WGD predicted from the other two | 337 | $T_{\text{Loo}} > 0$, $P \leq 0.01$ | $T_{\text{Loo}} = 0.221$,<br>$P = 1/100,001$ | Met |
| Apple–pear WGD from reference score | 373 | $\rho > 0$, $P \leq 0.01$, lower bound $> 0$; $\rho \geq 0.10$ | $\rho = 0.169$ (0.069–0.262),<br>$P = 0.00047$ | Met |
| TGD from reference score | 151 | $\rho > 0$, $P \leq 0.05$, lower bound $> 0$; then $\rho \geq 0.10$, $P \leq 0.01$ | $\rho = 0.107$ (–0.050 to 0.260),<br>$P = 0.094$ | Not met |
| Budding-yeast WGD from reference score | 186 | As for TGD | $\rho = 0.280$ (0.131–0.418),<br>$P = 0.00009$ | Met |
| TGD and snail WGDs (no plant data) | 146 | $r \geq 0.25$, $P \leq 0.01$, lower bound $> 0$ | $r = 0.226$ (0.061–0.385),<br>$P = 0.0033$ | Positive; below effect threshold |
| Snail WGD from reference score | 25 | $r > 0$, $P \leq 0.05$, lower bound $> 0$ | $r = 0.033$ (–0.303 to 0.340),<br>$P = 0.44$ | Not met (low power) |

The association is graded rather than threshold-like. The probability of retention after a target WGD increased steadily with the number of descendant genomes of another WGD that retained both ohnologs (Fig. 2C, D). Among the 337 HOGs present in all three reference WGDs, the probability of retention after any one of them was 23–26% when neither of the other two had retained the gene, 42–45% when one had and 48–51% when both had (Fig. 2E). Consequently, 28 HOGs were retained after all three WGDs, against 16.2 expected if the events were independent given their marginal retention rates (1.72-fold; permutation *P* = 0.0008), and 111 were returned to single copy after all three, against 86.7 expected (1.28-fold; *P* = 1.0 *×* 10*^−^*^4^; Fig. 2B). Repeated retention and repeated loss are therefore both more common than chance.

### About 30% of gene-level variation in retention propensity is shared across WGDs

Pairwise correlations of about 0.2 cannot be read directly as the strength of a shared tendency, because each WGD is observed through only four to six genomes and because much of the variation in retention is specific to each event. To separate these sources we fitted a liability-threshold model, the standard model of quantitative genetics for binary traits (Falconer 1965). In this model an unobserved, continuous retention propensity (the liability) determines whether a given descendant genome retains both ohnologs, and the liability is the sum of three parts: a component of the ancestral gene shared by all WGDs, a component of the gene specific to one WGD and shared by all descendants of that WGD, and a component specific to one descendant genome, including chance losses on terminal branches (Methods). Descendants of the same WGD share the first two components, whereas descendants of different WGDs share only the first, so the model is identified by comparing agreement within and between WGDs. Agreement was measured by tetrachoric correlations, which estimate the correlation of the underlying liabilities from two binary outcomes (Pearson 1900). Within-WGD correlations were high (0.43–0.86) and between-WGD correlations lower but consistently positive (Fig. 3A). In simulations with the same structure, the fitting procedure recovered known repeatabilities of 0.17, 0.31 and 0.43 as 0.19, 0.31 and 0.43 on average (Supplemental Note S3).

**Figure 3.**
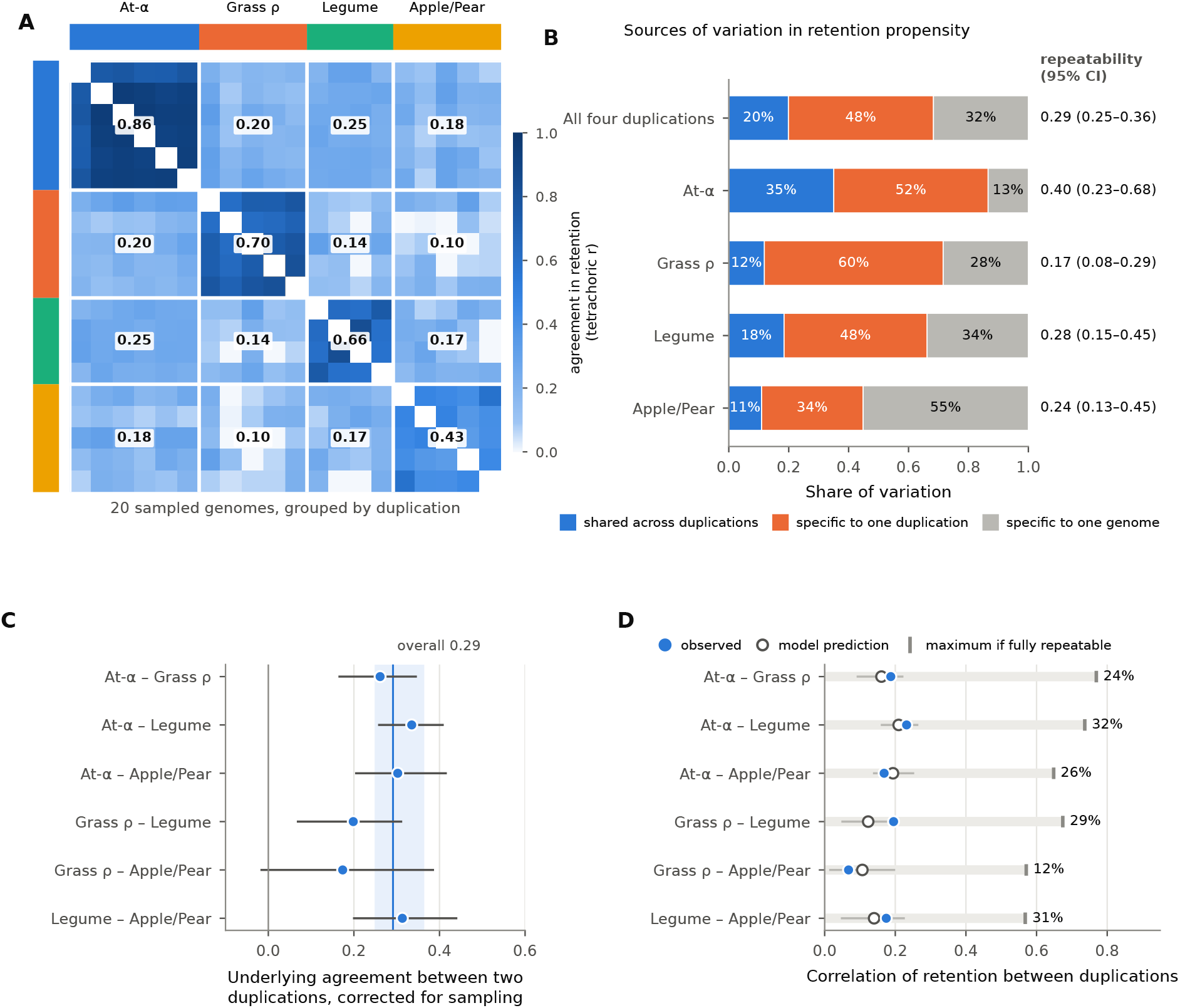
About 30% of gene-level variation in retention propensity is shared across WGDs. **A** Tetrachoric correlations of retention between all pairs of the 20 plant descendant genomes; numbers are block means. **B** Liability variance partitioned by the liability-threshold model into components shared across WGDs, specific to one WGD and specific to one genome, for all four WGDs jointly and per WGD; right, repeatability (shared component as a fraction of gene-level variance) with 95% bootstrap intervals. **C** Latent correlation for each pair of WGDs, disattenuated for within-WGD agreement; the band shows the overall repeatability and its interval. **D** Observed correlations of retention percentiles (blue), correlations predicted by the fitted model (open circles) and the maximum attainable if all gene-level variation were shared (grey ticks); percentages give the observed value as a share of the maximum.

Across the four plant WGDs and their 20 descendant genomes, the shared component accounted for 20% (95% CI 17–25%) of liability variance, the WGD-specific component for 48% (44–52%) and the descendant-specific component for 32% (30–33%) (Fig. 3B; loadings of each genome in Supplemental Fig. S3). We define repeatability as the shared component divided by the total gene-level (shared plus WGD-specific) variance (Nakagawa and Schielzeth 2010); it is the expected correlation between the propensities of the same ancestral gene in two independent WGDs. Repeatability was 29% (95% CI 25–36%) overall and ranged from 17% (grass *ρ*) to 40% (At-*α*) among events (Fig. 3B). Pairwise latent correlations, obtained by disattenuating the mean between-WGD tetrachoric correlation for within-WGD agreement (Spearman 1904), ranged from 0.17 to 0.34; the 95% interval excluded zero for five of the six pairs, the exception being grass *ρ*–apple–pear (0.17, −0.02 to 0.39; Fig. 3C).

The model also accounts for the size of the observed correlations. Simulating data from the fitted model with all gene-level variation assigned to the shared component gives the largest correlation the present sampling allows: 0.67–0.77 for the reference pairs. The observed correlations are 24–32% of this maximum, and they lie within the 95% intervals of the correlations predicted by the fitted model for five of the six plant pairs and at the upper limit for the sixth (grass *ρ*–legume; Fig. 3D). Pairwise correlations near 0.2 are thus the expected signature of a shared component of about 30% observed through a few genomes per event, not evidence of a weak or unreliable association.

### Gene properties account for about one-third of the association between WGDs

A shared component could arise because the same classes of genes are favoured after every WGD. To test this we described each ancestral gene by 76 features in seven classes: function (45 Gene Ontology [GO] slim categories and annotation depth), gene-family size and duplication history, protein–protein interactions, sequence conservation and gene age, essentiality (lethal and embryo-defective mutant phenotypes), dosage sensitivity (expression responses to aneuploidy and polyploidy, and membership of conserved protein complexes), and expression level, breadth and protein abundance (Methods; Supplemental Note S4). Individually, these features behaved as expected from previous work: transcription factors, kinases, signalling proteins, conserved complex members and highly connected proteins were preferentially retained, and plastid-targeted proteins preferentially lost (Supplemental Table S3).

We then removed from each WGD’s retention the part predictable from all features and recomputed the association between WGDs. Under the model above, the association between WGDs arises only from the shared component, so the fraction of the association removed by adjustment estimates the fraction of the shared component captured by the features. Three estimators were used because each has a different bias: in-sample least squares removes everything linearly predictable, including chance alignment, which we corrected by subtracting the fraction removed after permuting feature rows; cross-fitting, in which each gene’s adjustment comes from a model fitted to other genes, removes only effects that generalize (Chernozhukov et al. 2018); and a ridge model added 488 individual GO terms (Hoerl and Kennard 1970). The pooled correlation among reference WGDs fell from 0.208 (95% CI 0.174–0.241) to 0.139 after cross-fitted adjustment (averaged over five fold assignments), to 0.131 (0.092–0.170) after in-sample adjustment and to 0.152 with the GO terms added (Fig. 4B; Supplemental Table S6). The features thus accounted for 33% (cross-fitted) to 40% (in-sample, chance-corrected) of the association between WGDs. Function contributed most (14%), followed by protein interactions (8%), family size (6%), dosage sensitivity (5%), conservation and age (4%), expression (4%) and essentiality (1%) (Fig. 4A; Supplemental Fig. S4). Because measured features are imperfect proxies of the underlying properties, all three estimators under-adjust, and the residual association is an upper bound on the component independent of those properties.

**Figure 4.**
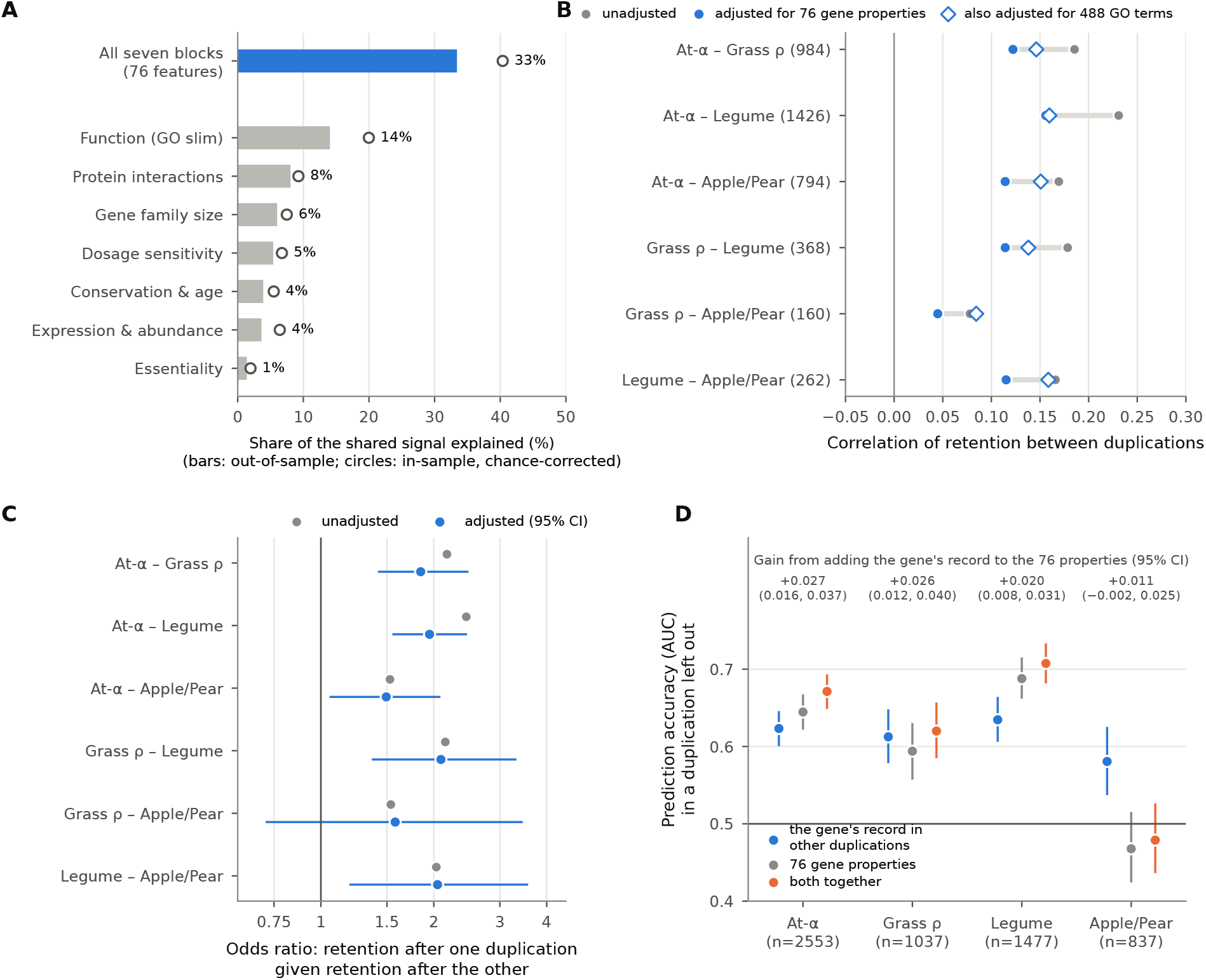
Gene properties explain part of the shared component, and retention history adds predictive information. **A** Percentage of the association among reference WGDs removed by adjustment for each feature class and for all 76 features (bars, cross-fitted; circles, in-sample after subtracting the chance contribution). **B** Correlation of retention percentiles before (grey) and after (blue) adjustment for all features, and after additional adjustment for 488 GO terms (open diamonds); HOG numbers in parentheses. **C** Odds ratios before (grey) and after (blue, 95% interval) adjustment by cross-fitted risk scores. **D** AUC for retention in a held-out plant WGD predicted from retention history in the other plant WGDs, from the 76 features, or from both; top, gain in AUC from adding history to the features (95% interval).

On the odds scale, after adjustment by cross-fitted feature-based risk scores for both WGDs, retention after one reference WGD remained associated with approximately twofold higher odds of retention after another: adjusted OR 1.85 (95% CI 1.42–2.47) for At-*α*–grass *ρ*, 1.95 (1.55–2.45) for At-*α*–legume and 2.09 (1.37–3.32) for grass *ρ*–legume, compared with unadjusted values of 2.17, 2.44 and 2.15 in the same HOGs, which are those with feature data (Fig. 4C). Most features were measured in *Arabidopsis thaliana*, a descendant of At-*α*; the grass *ρ*–legume comparison, which involves neither *A. thaliana* nor its WGD, was nonetheless as robust to adjustment as the others. For apple–pear, adjusted ORs were 2.05 (1.19–3.57) with the legume WGD and 1.49 (1.06–2.08) with At-*α*.

Two further analyses address properties that are shared within gene families or were tested individually. First, among HOGs of the same OMA gene family, after centring each WGD’s retention on its family mean, retention remained correlated between reference WGDs (pooled *r* = 0.132, within-family permutation *P* = 0.034; 331 HOGs; Supplemental Fig. S5); everything that members of a family have in common, including family size and most functional annotation, is removed by this comparison. Second, earlier in the study we tested individual mechanistic proxies, each under rules fixed before its own outcome was examined (Supplemental Note S7; Supplemental Fig. S6). The yeast fitness cost of an extra gene copy, transcriptional responses to trisomy and conserved sequence constraint were not associated with retention, and conserved expression magnitude was associated with retention but did not measurably reduce the association between WGDs. A prespecified multivariate model of these properties, fitted on two reference WGDs, did not outperform single properties on the third.

### A gene’s retention history adds predictive information beyond its annotated properties

If the shared component is real, retention in some WGDs should predict retention in another beyond what annotated properties predict. For each plant WGD we compared three predictors of retention in that WGD: the mean retention percentile of the same HOG in the other plant WGDs, which involves no fitted parameters; a cross-fitted ridge-logistic model of the 76 features; and a model combining both (Fig. 4D; Supplemental Table S7). The history score alone reached AUCs of 0.58–0.63, similar to the feature model for grass *ρ* (0.61 versus 0.59) and At-*α* (0.62 versus 0.64; 95% CI of the difference −0.051 to 0.007) and lower for the legume WGD (0.63 versus 0.69). For apple–pear the *A. thaliana*-based feature model did not predict retention (AUC 0.47), whereas the history score did (0.58, 0.54–0.63). Adding the history score to the features improved prediction in all three reference WGDs, by 0.027 (95% CI 0.016–0.037) for At-*α*, 0.026 (0.012–0.040) for grass *ρ* and 0.020 (0.008–0.031) for the legume WGD. A gene’s retention history therefore carries information that its measured properties do not. These AUCs are modest: at the start of the study, thresholds were declared for this type of leave-one-WGD-out prediction (mean AUC *≥* 0.65 to pass), and the mean of 0.61 would be classed as weak under them, although the executed protocol replaced these thresholds with the rank-based tests in Table 2 (Supplemental Note S6). We therefore interpret the history score as carrying transferable information rather than as an accurate predictor of individual genes.

### Prespecified predictions succeed in the apple–pear and budding-yeast WGDs

A more demanding test of a shared component is prediction of WGDs that played no part in building the predictor, with the prediction fixed before the outcome is seen. The study was organized around such tests (Table 2; Methods). For each target WGD, the orthology map, the eligible HOGs, the predictor and the decision rules were fixed first; the resulting prediction table was hashed (SHA-256) and committed to a version-controlled repository; and only then was the target’s retention outcome constructed or opened and joined to the prediction. The predictor was the same for every target: the unweighted mean of a HOG’s retention percentiles in the available reference WGDs (at least two), with no fitted weights. The apple–pear, TGD and yeast tests were run in an order fixed in advance on 28–29 August 2026; the animal tests were specified afterwards, on 3 September, each frozen before its own association was computed. A test that failed its criterion was reported as failed and could not be rescued by secondary analyses. The freeze records, hashes and commit identifiers are listed in Supplemental Table S11 and are included in the reproducibility package. There was no external preregistration, and the effect-size, variance-component, covariate, aggregation and within-genome analyses reported in the preceding and following sections were added later and are estimates rather than confirmatory tests.

In the apple–pear WGD, the prespecified score was correlated with retention among the 373 eligible HOGs (Spearman *ρ* = 0.169, 95% CI 0.069–0.262; one-sided permutation *P* = 0.00047), meeting both prespecified criteria. Expressed on an interpretable scale, 35% of HOGs in the top quartile of the score were completely retained, against 21% in the bottom quartile (ratio 1.67, 95% CI 1.05–2.80; AUC 0.56, 0.50–0.62; Fig. 5A).

**Figure 5.**
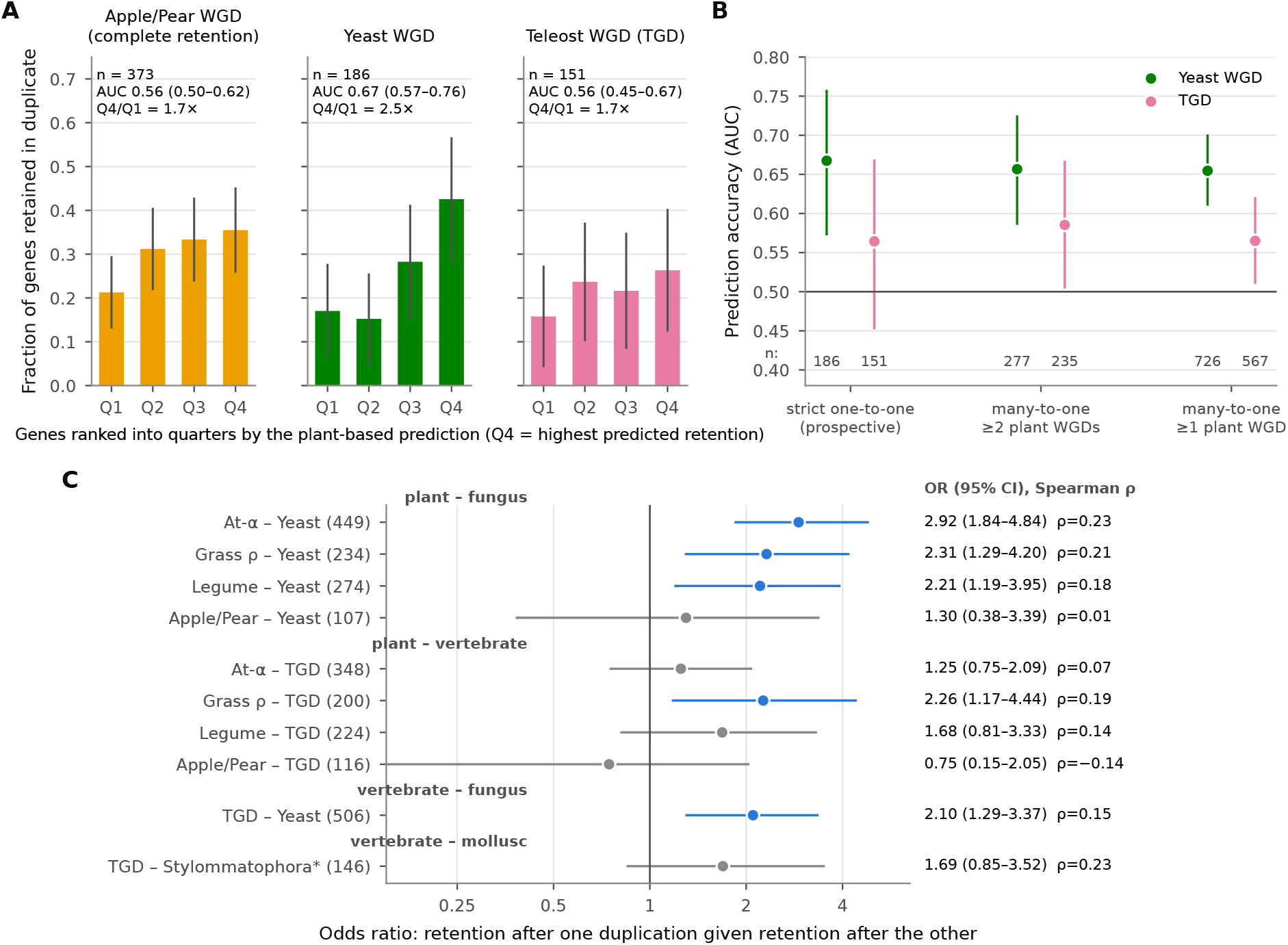
Prespecified predictions from angiosperm WGDs succeed in independent WGDs, including the budding-yeast WGD. **A** Fraction of HOGs retained in three target WGDs by quartile of the prespecified plant score (95% bootstrap intervals). AUC, area under the ROC curve; Q4/Q1, ratio of top to bottom quartile. **B** AUC for the yeast and TGD WGDs in the prespecified strict one-to-one populations and in the enlarged many-to-one populations of the sensitivity analysis; numbers of HOGs along the bottom. **C** Odds ratios (95% intervals) for retention in pairs of WGDs from different kingdoms; blue, interval excludes 1; right, odds ratios and Spearman correlations. *Stylommatophoran retention: both ohnologs in at least two of three genomes.

The budding-yeast WGD lies on a lineage that last shared an ancestor with plants about 1.5 billion years ago (Kumar et al. 2022). The prespecified plant score was nonetheless correlated with yeast retention among the 186 eligible HOGs (*ρ* = 0.280, 0.131–0.418; *P* = 0.00009), meeting both criteria: 43% of top-quartile HOGs retained both ohnologs in at least one yeast genome, against 17% of bottom-quartile HOGs (ratio 2.50, 1.30–6.37; AUC 0.67, 0.57–0.76). The TGD test, conducted before the yeast test, did not meet its criterion: the association was positive but imprecise among 151 HOGs (*ρ* = 0.107, −0.050 to 0.260; *P* = 0.094; top versus bottom quartile 26% versus 16%). The yeast WGD had been ranked as the next distant target before the TGD test, and its test used the same predictor construction and decision rules; the TGD result did not change any of them.

The deep prespecified populations were small because they required a one-to-one lift from mesangiosperm to eukaryotic HOGs. In a later sensitivity analysis, we averaged the scores of all angiosperm HOGs descending from the same eukaryotic HOG. Keeping the requirement of at least two reference WGDs enlarged the populations to 277 HOGs for yeast and 235 for the TGD (yeast AUC 0.66, 0.59–0.73; TGD AUC 0.59, 0.50–0.67, *P* = 0.019), and relaxing it to one reference WGD enlarged them to 726 and 567 (yeast AUC 0.65, 0.61–0.70, *P* = 0.0005, top-to-bottom quartile ratio 3.4; TGD AUC 0.57, 0.51–0.62, *P* = 0.010; Fig. 5B). The prespecified TGD result is therefore best explained by a smaller effect measured in a small sample, although this conclusion rests on an analysis that was not prespecified.

### Concordance extends across kingdoms

Direct pairwise comparisons at the eukaryotic level were consistent with these results (Fig. 5C; Supplemental Table S8). HOGs retained after each reference angiosperm WGD were more often retained after the yeast WGD: OR 2.92 (95% CI 1.84–4.84; *n* = 449; rank correlation permutation *P* = 1.0 *×* 10*^−^*^4^) for At-*α*, 2.31 (1.29–4.20; *n* = 234; *P* = 0.0008) for grass *ρ* and 2.21 (1.19–3.95; *n* = 274; *P* = 0.0018) for the legume WGD. Retention after the TGD and after the yeast WGD were also associated (OR 2.10, 1.29– 3.37; *n* = 506; *P* = 0.001), a comparison that uses no plant data. Associations between the angiosperm WGDs and the TGD were positive but less consistent (OR 1.25–2.26), with the interval excluding 1 only for grass *ρ*. The apple–pear WGD, whose complete-retention outcome mainly reflects recent losses among closely related species, was not associated with the yeast or TGD outcomes. Latent correlations estimated as for the plant WGDs were 0.32–0.41 between the reference angiosperm WGDs and the yeast WGD and 0.31 (0.08–0.45) between the TGD and yeast WGDs, comparable to those among angiosperm WGDs (Supplemental Note S5, Supplemental Fig. S7). HOGs that can be followed from angiosperms to fungi and animals are older and more conserved than average, so these estimates apply to that subset.

Within animals, retention after the TGD and after the stylommatophoran WGD was positively correlated among 146 bilaterian HOGs (*r* = 0.226, 95% CI 0.061–0.385; *P* = 0.0033). This prespecified test passed its inferential criteria but fell short of its prespecified strong-effect threshold (*r ≥* 0.25). A strict comparison between the plant score and snail retention could be made for only 25 HOGs and was unresolved (*r* = 0.033, −0.303 to 0.340; *P* = 0.44); with this sample size only correlations above about 0.5 were detectable.

### The findings are robust to orthology rules, descendant sampling and dependence

Because the analysed sets shrink with taxonomic depth, comparisons share genes and events, and WGDs are represented by different numbers of genomes, we tested each of these features directly (Supplemental Note S2, Supplemental Figs. S1–S2, Supplemental Tables S4–S5). Pillars excluded at the mesangiosperm level, because they lacked sufficient exact matches, because their two retained ohnologs matched different HOGs, or because they shared a HOG with another pillar of the same event, were retained more often than analysed pillars (for example 54% versus 36% for At-*α*; Supplemental Table S4). Exclusions that depend on each WGD’s outcome separately, such as those for within-event multiplicity and within-species conflict, multiply the joint distribution of outcomes by factors that cancel from the odds ratio, so they cannot create an association between WGDs (Methods); for the remaining exclusions we relied on the alternative rules below. Empirically, requiring exact matches from at least three species, retaining multiplicity-affected HOGs by averaging their pillars, restricting to HOGs with a one-to-one eukaryotic lift, or using a different taxonomic level for mapping all gave pooled correlations of 0.182–0.230 and ORs of 1.76–2.81, against 0.210 for the primary map.

To equalize sampling, we recomputed retention occupancy for every combination of four descendant genomes per reference WGD (75 combinations): the pooled correlation had a median of 0.202 (range 0.183–0.216) and pairwise ORs ranged from 1.92 to 2.60. To account for dependence, we resampled HOGs jointly across all pairs, estimated the covariance of the pairwise estimates and pooled them by generalized least squares, which gave 0.210 (95% CI 0.172–0.248), nearly identical to the pooled estimate that ignores dependence; the pairwise estimates were nearly uncorrelated (mean correlation 0.048) because each pair is dominated by HOGs absent from the others. Omitting any one plant WGD gave pooled values of 0.161–0.210. Finally, the snail reconstruction was checked by two independent phylogenetic analyses: in a prespecified panel of 300 loci analysed with maximum-likelihood trees, 8 of 213 informative trees (3.8%) conflicted with the inferred assignment of genes to duplicate tracks (Supplemental Note S1).

### Within-genome tests: yeast genomes lose protein-complex partners together

Gene properties and retention histories describe which genes tend to be retained, but on their own they cannot separate a cause from a correlate, because any measured property may stand in for an unmeasured one. We therefore used a design in which the gene itself is held constant. Within one WGD, the same ancestral gene is retained in some descendant genomes and returned to single copy in others. If dosage balance shapes retention, a genome that has lost a gene’s stoichiometric partners should more often lose that gene as well, because the partner loss removes the imbalance that losing the second copy would otherwise create. We modelled each gene’s two-copy state in each genome on the fraction of its complex partners retained in that genome, with fixed effects for gene and for genome (Fig. 6A; Methods). The gene effects absorb every property of the gene, including function, expression, family size and constraint, and the genome effects absorb each genome’s overall rate of loss. Partners lying within 20 genes in the ancestral order were excluded, so that a single deletion could not remove both, and significance was assessed against partner sets permuted among complex members.

**Figure 6.**
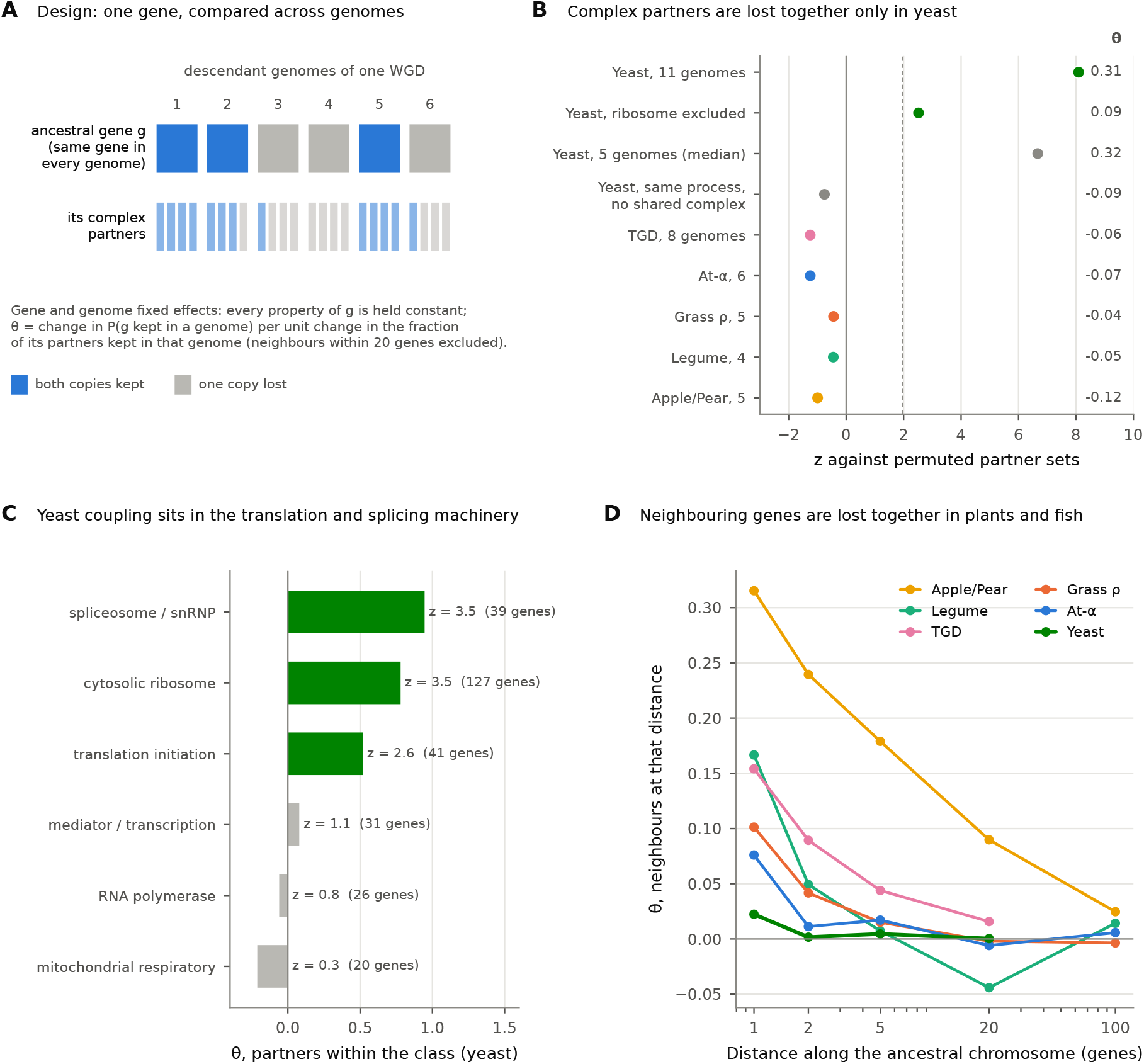
Within-genome tests separate selection on dosage balance from mutational loss. **A** Design. The same ancestral gene is compared across the genomes descended from one WGD; gene and genome fixed effects hold every property of the gene and each genome’s overall loss rate constant, and *θ* is the change in the probability that a genome keeps both copies of the gene per unit change in the fraction of its complex partners kept in that genome (partners within 20 genes in the ancestral order excluded). **B** Coupling between complex partners for each WGD and for yeast controls, as *z* against permuted partner sets; *θ* at right. **C** Yeast coupling within named complex classes, with *z* against random member sets of the same size; classes in which almost no genes vary across genomes are omitted. **D** Coupling between a gene and the genes at a given distance along the ancestral chromosome, for all POInT pillars of each WGD.

In the 11 genomes descended from the yeast WGD, partner loss and gene loss were strongly coupled (Fig. 6B; Supplemental Table S12). A genome that retained none of a gene’s partners was 31 percentage points less likely to retain that gene than a genome that retained all of them (*θ* = 0.31, 95% CI 0.22–0.41; permuted-partner null −0.005 *±* 0.040; *z* = 8.1; *P* = 0.0005, the smallest attainable; 697 genes, 5,881 partner pairs). The coupling was unchanged by the width of the neighbour exclusion (*z* = 7.8–8.4 for 0 to 100 genes), by omitting any one genome (*θ* = 0.27–0.33, *z* = 6.5–9.4) and by the source of complex annotations (*z* = 5.0 with CORUM alone to 9.1 with CYC2008 alone) (Supplemental Table S13).

The coupling has the specificity that dosage balance predicts. It was strongest in large stoichiometric assemblies of the translation and splicing machinery: the cytosolic ribosome (*θ* = 0.78, *z* = 3.5), the spliceosome (*θ* = 0.95, *z* = 3.5) and translation-initiation complexes (*θ* = 0.52, *z* = 2.6) (Fig. 6C). Removing any one complex class other than the ribosome left the coupling intact (*θ* = 0.31–0.33, *z* = 8.0– 8.9); without ribosomal proteins it was weaker but still present (*θ* = 0.09, *z* = 2.5, *P* = 0.008; bootstrap interval −0.01 to 0.19). It was confined to physical partners. Genes that share a biological process but no complex showed no coupling (*θ* = −0.09, *z* = −0.8), whereas partners that share both a complex and a process showed the strongest coupling outside the ribosome (*θ* = 0.22, *z* = 3.1). A change in a lineage’s demand for a whole pathway would couple same-process genes too; specificity to physical partners points instead to stoichiometry. Coupled losses also occurred on recent branches. In four sister-species pairs, when a gene had been lost in one sister only, its partners had been lost in the same sister in 68% of cases, against 53 *±* 7% with permuted partners (*P* = 0.027). Whether partner loss precedes gene loss could not be resolved with these data (*P* = 0.221). Genes whose extra copy is costly in a laboratory gain-of-dose assay (MoBY-CEN) (Morrill and Amon 2019) tended to be more strongly coupled than genes whose extra copy is cheap (*θ* = 0.55 versus 0.29; one-sided *P* = 0.087).

No such coupling was detected after the plant WGDs or the TGD (*z* = −1.8 to 1.4; pooled *z* = −1.96; Fig. 6B). This absence is informative. Reduced to 4, 5 or 6 genomes, the yeast data still detected the coupling in 92%, 95% and 100% of random subsets, and the upper 95% bound of *θ* in every plant and fish WGD (at most 0.10) lies below the lower bound in yeast (0.22).

### Neighbouring genes are lost together in plants and fishes, but not repeatedly

The same design applied to neighbours in the ancestral gene order revealed a different process (Fig. 6D; Supplemental Table S14). After the plant WGDs and the TGD, a genome that had lost the adjacent gene was 8–32 percentage points more likely to have lost the focal gene (*θ* = 0.08–0.32, *z* = 7.5–38.2), and the effect decayed within two to five genes after the older plant WGDs and within about 20 genes after the TGD. This is the signature of short deletions that remove one or a few adjacent genes at once (Woodhouse et al. 2010). In yeast genomes the effect was nearly absent (*θ* = 0.02). In the young apple–pear WGD it extended over tens of genes, partly because of one genome (*θ* = 0.17 without *Pyrus bretschneideri*).

This positional process decides which genome loses a gene but does not explain why the same genes are retained after independent WGDs. Neighbours explained at most 4% of a gene’s retention level within a WGD. The retention of a gene’s neighbours after one WGD did not predict its own retention after another (*ρ* = −0.11 to 0.00), and removing each WGD’s regional component left most of the association between WGDs in place (for example 0.23 to 0.20 for At-*α* and legume). The repeatable part is therefore a property of the gene, not of its position. Positional losses add lineage-specific variation instead, which helps to explain why repeatability is about 30% rather than higher.

### Constraint before duplication marks the shared propensity across kingdoms

To ask which properties carry the repeatable part, we divided each plant gene’s propensity into its shared and WGD-specific components using the fitted liability model (Methods). Every measured property associated more strongly with the shared than with the WGD-specific component, but none exceeded *|ρ|* = 0.17 (Fig. 7A; Supplemental Table S15). Expression level measured in species outside the duplicated lineages was the strongest (shared *ρ* = 0.17, WGD-specific *ρ* = 0.04; 316 genes), and its association persisted after adjustment for sequence constraint (partial *ρ* = 0.14). Expression is therefore linked to the repeatable propensity, but it accounts for only about 3% of its variance, which explains why the earlier prespecified test detected no attenuation of replay (Supplemental Note S7).

**Figure 7.**
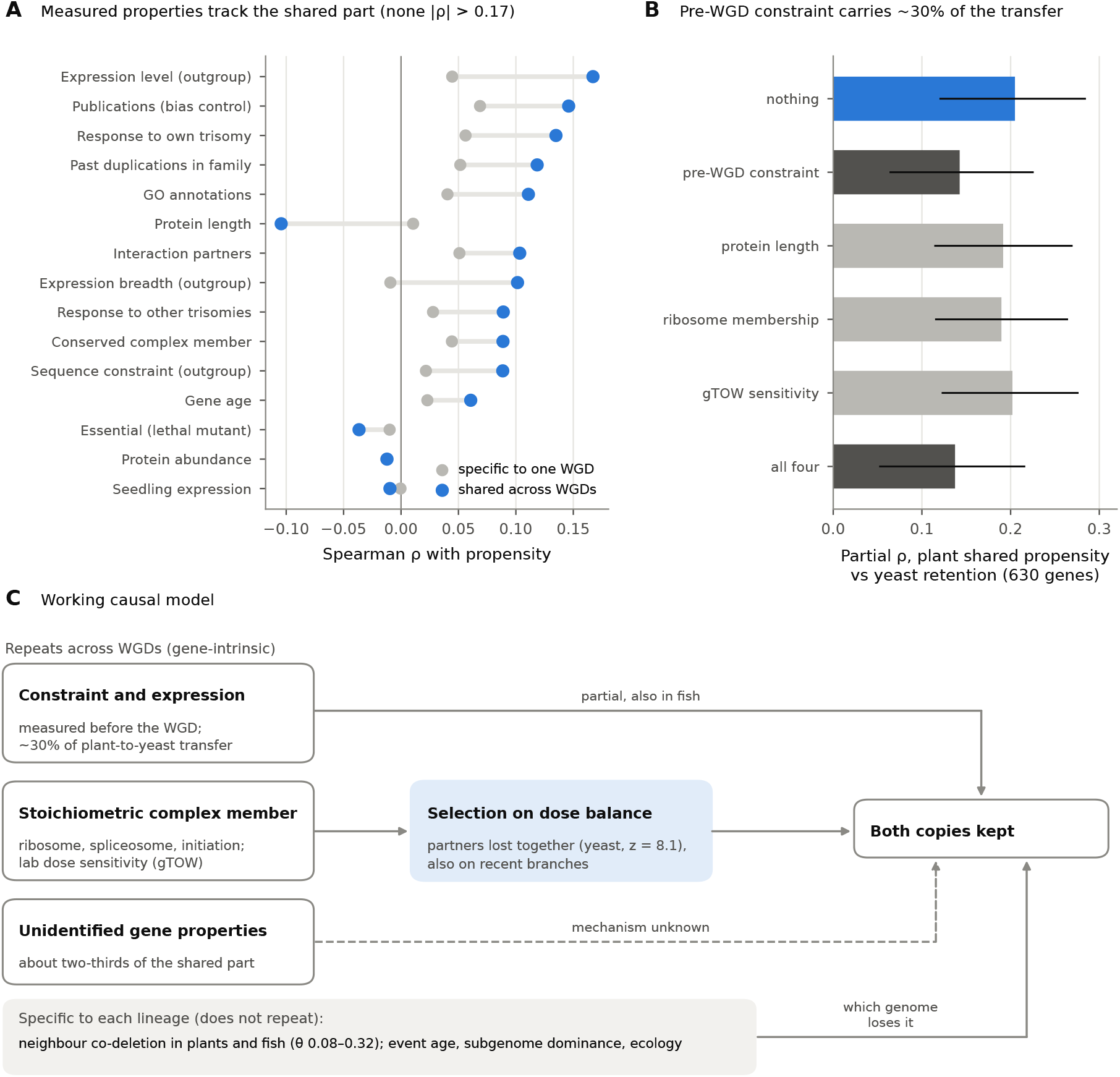
What carries the shared propensity. **A** Spearman correlation of gene features with each plant gene’s shared propensity (blue) and WGD-specific propensity (grey), estimated from the liability-threshold model for genes present in at least two WGDs. **B** Partial correlation between the plant shared propensity and retention after the yeast WGD for 630 genes, conditioning on nothing or on pre-duplication properties (95% bootstrap intervals). **C** Working causal model. Solid arrows are supported by the analyses in this study; the dashed arrow marks the unexplained part of the shared propensity.

Properties measured before duplication cannot be consequences of retention. Protein-sequence constraint estimated in outgroups that did not undergo the WGD predicted retention after the yeast WGD (*ρ* = 0.26, 1,163 genes, *P* = 2.1 *×* 10*^−^*^19^; *ρ* = 0.17 without ribosomal proteins) and after the TGD (*ρ* = 0.17, 615 genes, *P* = 1.5*×*10*^−^*^5^). Pre-duplication protein length did not (yeast *ρ* = 0.00 without ribosomal proteins; TGD *ρ* = −0.04), which argues against a neutral model in which longer genes are lost faster because they present a larger mutational target. In a laboratory test of dose tolerance, yeast genes that tolerate fewer extra copies (gTOW copy-number limit) (Makanae et al. 2013) were more often retained after the yeast WGD (*ρ* = 0.16, 1,380 genes, *P* = 3.6 *×* 10*^−^*^9^; *ρ* = 0.14 without ribosomal proteins).

Constraint also carried part of the cross-kingdom signal. Among 630 genes shared by the plant and yeast analyses, the plant shared propensity predicted yeast retention (partial *ρ* = 0.205, 95% CI 0.12–0.28). Holding pre-duplication constraint fixed reduced this to 0.143 (0.06–0.23), a reduction of 30%, whereas holding protein length, ribosomal membership or gTOW sensitivity fixed changed it by less than 0.02 (Fig. 7B). Constraint itself was shared: constraint measured in yeast outgroups correlated with the plant shared propensity (*ρ* = 0.249). Finally, the shared propensities of plant complex partners were correlated (22,485 partner pairs, *z* = 5.6 against permuted complexes), so part of the propensity belongs to the complex rather than to the single gene. Figure 7C summarizes how these results fit together.

## Discussion

Retention of ohnologs after independent WGDs is partly repeatable at the level of individual ancestral genes. A gene retained after one angiosperm WGD was 1.67–1.73 times as likely to be retained after another, genes retained after all three reference WGDs were 1.72-fold over-represented, and predictions fixed before the outcomes were examined succeeded in the apple–pear WGD and in the budding-yeast WGD. Within-genome comparisons, which hold each gene fixed, connect this repeatability to a mechanism: in yeast, genomes lose stoichiometric partners together. These effects are moderate for any single gene and do not make the fate of an individual duplicate predictable with certainty. They are, however, consistent in direction across comparisons, robust to analytical choices and, most importantly, observed in WGDs that contributed nothing to the prediction.

The variance decomposition specifies the kind of repeatability involved. About 30% of gene-level variation in retention propensity is shared across WGDs and about 70% is specific to each event. In the terminology of quantitative genetics, repeatability measures the fraction of variation attributable to a persistent property of the unit, here the ancestral gene, as opposed to the particular occasion, here the WGD (Nakagawa and Schielzeth 2010). A large event-specific component is expected. WGDs differ in age, in whether they arose by autopolyploidy or allopolyploidy, in subgenome dominance and biased fractionation (Thomas et al. 2006; Schnable et al. 2011; Emery et al. 2018; Cheng et al. 2018), and in the ecological and demographic history of the surviving lineage, and some losses are chance events in a single descendant. Against this background a shared component of almost a third is substantial: the identity of the gene contributes measurably to its fate, even though the history of the event contributes more. The decomposition also resolves an apparent inconsistency in the literature, in which strong functional biases in retained ohnologs coexist with modest gene-level correlations between events; the correlations are attenuated by sparse descendant sampling and by the event-specific component, and reach 24–32% of their attainable maximum.

Measured gene properties account for about one-third of the association between WGDs, and hence of the shared component. Function and protein interactions contribute most, consistent with dosage balance as a major determinant of biased retention (Maere et al. 2005; Papp et al. 2003; Birchler and Veitia 2007; Tasdighian et al. 2017). The remainder persisted after adjustment for 76 features and nearly 500 GO terms, persisted within gene families, and improved prediction beyond all features. Part of this remainder may reflect properties that we measured incompletely, because most features come from *A. thaliana* and from imperfect annotations, and all adjustment estimators under-adjust for the latent properties they proxy. Part probably reflects properties that single-gene annotations represent poorly, such as the stoichiometry of a protein within its complexes and whether its partners were also retained. The within-genome tests show that this second possibility is real. In yeast, a genome that lost a gene’s physical partners was 31 percentage points less likely to retain that gene, an effect that no property of the gene can produce because the gene is the same in every comparison. The coupling was specific to physical partners, strongest in the ribosome, spliceosome and translation-initiation machinery, present on recent branches and stronger for genes whose extra copy is costly. This is the pattern expected if selection acts on the balance of whole complexes, the central claim of the dosage-balance hypothesis (Birchler and Veitia 2007; Freeling 2009; Conant et al. 2014), and it explains why complex membership measured gene by gene captures so little: what matters is whether the partners are also present in two copies. The prespecified tests of single-gene proxies of dosage sensitivity reached the same conclusion from the other side: none accounted for the shared component.

Coupled losses of partners were not detected in plants or fishes, although the test had the power to detect a yeast-sized effect. Instead, losses in these lineages were coupled along the chromosome, which is the signature of short deletions removing adjacent genes (Woodhouse et al. 2010); similar clustered losses have been observed within decades of the origin of a natural allopolyploid (Buggs et al. 2012). Yeast genomes are compact, and their duplicates were lost largely one gene at a time, so that selection on each loss can be resolved. In the larger plant and fish genomes, many losses remove several genes at once, and the copies that remain may be buffered by paralogs from older duplications; both would hide partner coupling. The positional process is lineage specific. It decides which genome loses a gene but does not recur between WGDs, and it is one reason why repeatability is about 30% rather than higher. Expression and constraint occupy a different place in this picture. Both mark the repeatable part rather than the WGD-specific part, and constraint measured before duplication accounted for about a third of the plant-to-yeast transfer. Genes under strong purifying selection before a WGD are those whose dose and function are most constrained, and they are the genes whose duplicates are most often retained across kingdoms. Expression is a real but small contributor. It explains about 3% of the shared propensity, which is why it failed the prespecified test of whether it accounts for replay.

The cross-kingdom results indicate that part of the shared component is ancient. Plant WGDs predicted retention after the yeast WGD at least as well as they predicted the apple–pear WGD, and the TGD and yeast WGDs were concordant without any plant data. Many of the HOGs that can be followed across kingdoms encode core cellular machinery that has retained similar roles since the last common ancestor of eukaryotes (Davis and Petrov 2004), which would allow the same dosage or network constraints to act after WGDs in different kingdoms. The weaker prespecified TGD result fits the long and complex history that followed the teleost duplication, including extended rediploidization and lineage-specific losses (Brunet et al. 2006; Sémon and Wolfe 2007; Inoue et al. 2015; Conant 2020; Qi et al. 2024), but we did not estimate formal differences between targets, and the data do not support a simple decline of predictability with phylogenetic distance.

The findings also have practical implications. For a newly sequenced polyploid, including the many polyploid crops, the retention history of an ancestral gene across other WGDs provides an annotation-independent prior for whether its ohnologs are likely to be retained, and this prior adds information to functional annotation. Conversely, the large event-specific component cautions against expecting one WGD to mirror another closely, and identifies the history of each event as the larger source of variation to be explained.

Our study has limitations. Reconstructions record only loci that remain detectable, so ancestral genes lost from every descendant early after a WGD are under-represented. One-to-one mapping excludes some highly retained genes; this cannot create the association we report, as shown above, but restricts the population to which estimates apply. Most features were measured in *A. thaliana*, although conclusions held for comparisons not involving it. We analysed seven WGDs; more events, each with several descendant genomes, would sharpen the estimates and allow event-level predictors of repeatability to be tested. The snail reconstruction rests on three genomes. The within-genome coupling in yeast rests largely on ribosomal proteins, whose duplicate copies are kept nearly identical by gene conversion; robustness to omitting any genome and the specificity to physical partners argue against assembly artefacts, but locus-level checks would rule them out. Holding the gene fixed removes gene-level confounding but not a lineage-wide cause acting on a whole complex at once, such as selection on growth rate across all ribosomal proteins; reconstructing the order of losses on a dated tree, and testing the predicted alleviating genetic interactions between deletions of duplicate copies of partner genes in existing yeast interaction maps (Costanzo et al. 2016; Kuzmin et al. 2020), would separate these. Finally, the reference replay, apple–pear, yeast, TGD and animal tests were prespecified, whereas the effect-size, variance-component, covariate-adjustment, aggregation and within-genome analyses were added later and should be interpreted as estimates.

In conclusion, duplicate retention after whole-genome duplication is partly repeatable at the level of individual ancestral genes. About 30% of gene-level variation in retention propensity is shared across independent WGDs, and it predicts retention in prespecified tests extending to the budding-yeast WGD. Within-genome tests link this repeatability to dosage balance within protein complexes, and pre-duplication constraint accounts for part of it across kingdoms, whereas short deletions and lineage history decide where losses fall. A gene’s retention history is therefore an informative and portable, although partial, guide to its fate after genome duplication.

## Methods

### Whole-genome duplications and reconstructions

We used frozen, versioned reconstructions from POInT (Hao et al. 2022; Conant 2023) for the Brassicaceae At-*α* WGD (six descendant genomes), the Poaceae grass *ρ* WGD (five), the papilionoid legume WGD (four), the Maleae WGD (five), the budding-yeast WGD (eleven) and the TGD (eight) (Conant 2020). Release files, checksums and parsing audits are part of the reproducibility package, and descendant genomes are listed in Supplemental Table S2. For pillar *i* of event *e*, each descendant carries one or two WGD-derived copies; zero-copy cells are not represented in POInT and were treated as parsing failures.

The stylommatophoran WGD was reconstructed from the genomes of *Arion vulgaris*, *Lissachatina fulica* and *Bradybaena similaris*, with *Onchidella celtica* as unduplicated outgroup (Liu et al. 2021; Chen et al. 2022; McHale et al. 2025). All 12,570 protein-coding *O. celtica* genes on chromosomes 1–18 formed a loss-inclusive candidate set. Accepted two-track loci required significant enrichment of the outgroup chromosome (at least ten anchors per block), at least two additional compatible anchors within 20 genes, an accepted paralogon and correspondence of blocks across descendants; 618 high-confidence and 349 moderate-confidence loci were accepted. Snail proteins were mapped to OMA with a rule calibrated on a balanced panel of 5,400 held-out molluscan proteins (precision 0.9994, coverage 0.9957). Validation analyses are described in Supplemental Note S1.

### Retention outcomes

Retention occupancy was *y_ie_* = *k_ie_*/*m_e_*, where *k_ie_* is the number of descendants of event *e* carrying both ohnologs of pillar *i* and *m_e_* the event’s number of descendants. Occupancy describes the observed tip pattern and does not identify the branch on which a copy was lost. Event-wide percentiles *R_ie_* = (midrank(*y_ie_*) *−* 0.5)/*N_e_*, with average ranks for ties, were computed once over all *N_e_* mapped pillars of each event and were not recomputed within the intersections used for pairwise comparisons. The binary outcome was retention in at least one descendant for the At-*α*, grass *ρ*, legume, yeast and TGD events, and complete retention in all five genomes for the Maleae event (rationale in Results). For the stylommatophoran WGD, retention required both copies in at least two of three genomes.

### Orthology mapping

Translated POInT coding sequences were matched to OMA All.May2026 proteins by identical normalized peptide sequence within the same species (Altenhoff et al. 2024). For each pillar we recorded the HOGs supported by each exactly matched descendant. A pillar was eligible when at least two descendant species matched exactly, all matches supported a single HOG and no species supported several HOGs. Pillar– HOG relations were then required to be one-to-one within each event, and HOGs matched by several pillars of any reference event were excluded from primary analyses. Comparisons among plant WGDs used Mesangiospermae HOGs. Yeast and TGD pillars were mapped to Eukaryota HOGs with the same rules, and mesangiosperm HOGs were lifted to Eukaryota HOGs through the OMA hierarchy; the prespecified deep tests required a one-to-one lift. The TGD–snail comparison used Bilateria HOGs. None of the five Maleae genomes is represented in OMA, so Maleae pillars were mapped with DIAMOND 2.2.5 (Buchfink et al. 2021) against 84 mesangiosperm proteomes from the same OMA release, using a precision-first rule (support from at least two descendant species, unanimous HOG assignment, no within-species conflict, one-to-one relation) whose thresholds were calibrated on held-out exact assignments of the reference events before any Maleae outcome was examined.

### Association between pairs and triples of WGDs

For each pair of WGDs (*a, b*) we report, among shared HOGs, the probability of retention after *b* given retention or loss after *a*, the risk ratio, the odds ratio with a 0.5 continuity correction where a cell was empty, a one-sided Fisher’s exact test, the AUC of retention after *a* for discriminating retention after *b*, and the Pearson correlation *r_ab_*of event-wide percentiles. The prespecified global statistic was the (*n −* 3)-weighted Fisher mean over the reference pairs,

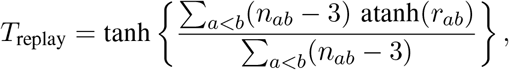

tested against 100,000 permutations that independently reassigned each event’s percentiles over fixed HOG slots, preserving the sparse pattern of shared HOGs; its interval came from 10,000 bootstrap resamples of HOGs that carried all event records of each HOG together. Intervals for pairwise statistics came from 10,000 bootstrap resamples of HOGs. Departure of joint retention from independence among the 337 HOGs present in all three reference WGDs was tested by permuting each WGD’s outcomes independently among these HOGs (10,000 permutations), which preserves each WGD’s retention rate.

### Liability-threshold model

For HOG *g*, WGD *e* and descendant genome *d*, the liability was modelled as

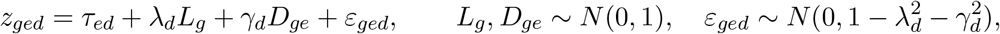

with both ohnologs present when *z_ged_ >* 0. The thresholds *τ_ed_* absorb differences in overall retention between descendants; *L_g_*is the gene’s propensity shared by all WGDs, *D_ge_* its WGD-specific propensity shared by all descendants of *e*, and *ε_ged_* everything specific to one descendant. This is a bifactor model (Holzinger and Swineford 1937) for binary indicators. Tetrachoric correlations (Pearson 1900) were computed for all 190 pairs of the 20 plant descendants on pairwise-complete HOGs with a 0.5 continuity correction, and the loadings *λ_d_, γ_d_ ∈* [0, 1) were estimated by weighted least squares fitting of the 190 correlations, weighting by the number of shared HOGs. Because the model implies *ρ_dd_′* = *λ_d_λ_d_′* + *γ_d_γ_d_′* within an event and *ρ_dd_′* = *λ_d_λ_d_′* between events, the shared loadings are identified only from correlations between descendants of different WGDs. Phylogenetic structure among descendants of one WGD is absorbed into *D_ge_* and *ε_ged_* and, if anything, leads to underestimation of repeatability. Repeatability was *̄λ*^2^/(*̄λ*^2^ + *̄γ*^2^), with bars denoting averages over descendants (Nakagawa and Schielzeth 2010). Pairwise latent correlations were the mean between-event tetrachoric correlation divided by the geometric mean of the two mean within-event correlations (Spearman 1904). The attainable maximum was obtained by simulating data from the fitted model with thresholds and total gene-level variance of each descendant unchanged but all gene-level variance assigned to *L_g_*. Intervals came from 1,000 bootstrap resamples of HOGs, refitting the model each time. Parameter recovery was checked by simulation (four events with six, five, four and five descendants; 4,000 HOGs; 20 replicates per scenario; Supplemental Note S3).

### Gene-property features and adjustment

*A. thaliana* genes were linked to Mesangiospermae HOGs through exact OMA matches (5,151 of 8,631 HOGs), and Eukaryota-level features were attached through the OMA hierarchy. Sources were: GO annotations, GO slim categories, mutant phenotypes and publication links from TAIR (Berardini et al. 2015; The Gene Ontology Consortium 2021); gene-family size and duplication history from OMA; gene age and protein-sequence constraint among single-copy orthologs of *Amborella*, grapevine, cacao and peach; multi-validated physical interactions from BioGRID 5.0 (Oughtred et al. 2021); essentiality from lethal, embryo-defective and gametophyte-lethal phenotypes and SeedGenes (Meinke et al. 2008; Lloyd and Meinke 2012); expression responses to aneuploidy and polyploidy (Hou et al. 2018); conserved membership of protein complexes from Complex Portal, CORUM and CYC2008 (Meldal et al. 2022; Giurgiu et al. 2019; Pu et al. 2009); and expression level, breadth and protein abundance in *A. thaliana* and three non-focal species (Wang et al. 2015; Fasoli et al. 2012; Flores-Tornero et al. 2020; Kulesza et al. 2024). Missing values were median-imputed with missingness indicators, and all 76 columns were standardized (column list in Supplemental Note S4).

Adjustment regressed each WGD’s retention percentile on the features and correlated the residuals between WGDs. (i) In-sample ordinary least squares removes everything linearly predictable, including chance alignment; we subtracted the mean fraction removed after permuting feature rows (200 permutations). (ii) Cross-fitted residuals were obtained from five-fold models fitted to other folds (Chernozhukov et al. 2018). (iii) A ridge model added 488 GO terms, with the penalty chosen by internal cross-validation (Hoerl and Kennard 1970). Block contributions were obtained by adjusting for each feature class separately. Adjusted ORs came from logistic regression of retention in one WGD on retention in the other plus cross-fitted ridge-logistic risk scores for both WGDs, because direct logistic adjustment for 76 features was unstable at these sample sizes. Within-family analyses grouped HOGs by OMA root family, kept families with at least two HOGs present in both WGDs, centred each WGD’s percentiles on family means, and assessed significance by permuting centred values within families (5,000 permutations).

### Prediction within plants

For each plant WGD, retention was predicted from the mean percentile of the same HOG in the other plant WGDs (no fitted parameters), from a cross-fitted ridge-logistic model of all features, and from both. Performance was measured by AUC with intervals from 2,000 bootstrap resamples, and the gain from adding the history score was estimated on the same resamples.

### Prespecified tests and freeze records

The study was conducted as a sequence of milestones, each with a written authorization that fixed its inputs, populations, statistics, random seeds and decision rules before the association it tested was computed. For the target tests, the procedure was: (i) fix the orthology map and eligible HOGs without reference to target outcomes; (ii) compute the predictor, the unweighted mean of available reference percentiles with at least two reference WGDs required; (iii) write the prediction table, record its SHA-256 hash and commit it; (iv) only then construct or open the target outcome, verify the committed hashes and join prediction to outcome once. The primary statistic was Spearman’s *ρ* with a one-sided test against 100,000 permutations of target labels within the frozen population and an interval from 10,000 paired bootstrap resamples. Decision rules are given in Table 2. The apple–pear test was followed by the distant targets in an order fixed in advance (TGD, then yeast); the TGD result was recorded as not meeting its criterion, and the yeast test used the same predictor construction and decision rules. The animal tests used outcomes constructed in earlier milestones but froze their populations and decision rules before the two outcomes were joined; their specification history, including earlier versions that stopped at sample-size gates before any association was computed, is given in Supplemental Note S6. Secondary analyses (source subsets, raw outcomes) were declared in advance and could not replace a failed primary result. Freeze records are summarized in Supplemental Table S11 (Supplemental Note S6), and the complete decision log is part of the reproducibility package. The study was not externally preregistered.

### Robustness analyses

Alternative orthology rules were: exact support from at least three species; averaging the pillars of HOGs with within-event multiplicity; restriction to HOGs with a one-to-one Eukaryota lift; and mapping at the Pentapetalae level for At-*α* and the legume WGD. Rarefaction recomputed occupancies from every combination of four descendants per reference WGD (15 *×* 5 *×* 1 = 75 combinations). Generalized least squares pooling used the covariance of Fisher-transformed pair estimates from 2,000 joint HOG resamples. Exclusion cannot create association when inclusion depends on each event’s outcome separately: if *R_a_, R_b_ ∈ {*0, 1*}* denote retention and *S* inclusion with *P* (*S* = 1 *| R_a_, R_b_*) = *f* (*R_a_*) *g*(*R_b_*), then among included HOGs *P* (*R_a_, R_b_ | S* = 1) ∝ *P* (*R_a_, R_b_*)*f* (*R_a_*)*g*(*R_b_*), and the factors *f* and *g* cancel from the odds ratio *P*_11_*P*_00_/(*P*_10_*P*_01_). Exclusions for within-event multiplicity and within-species conflict are of this form; they can still change the population to which estimates apply, which is why the relaxed rule is also reported.

### Within-genome coupling tests

For each WGD, *y_gd_* = 1 if descendant genome *d* retains both ohnologs of ancestral gene *g*. For a gene with at least one partner, *P_gd_* is the fraction of its partners retained in genome *d*. Partners were genes of the same WGD that share a stable heteromeric complex (CYC2008, CORUM or Complex Portal (Pu et al. 2009; Giurgiu et al. 2019; Meldal et al. 2022), mapped to Eukaryota HOGs; for plant WGDs also Gene Ontology complex terms), excluding paralogous lineages of one eukaryotic gene and genes within 20 pillars of *g* in the ancestral order. The coupling coefficient *θ* was estimated by least squares in *y_gd_* = *α_g_* +*β_d_* +*θP_gd_* +*e_gd_*, a linear probability model with gene and genome fixed effects computed by two-way demeaning of the balanced gene-by-genome panel. The null distribution came from permuting partner sets among complex-member genes (2,000 permutations), which keeps every gene’s own data and the neighbour exclusion; a second null replaced each member by a random gene of the same WGD with the same occupancy (500). Intervals for *θ* came from bootstrap resampling of focal genes. Robustness analyses varied the exclusion window, removed each genome, used each complex database alone, removed each complex class, restricted partners to named classes, contrasted physical partners with genes sharing a Gene Ontology biological process but no complex, and subsampled yeast genomes to 4–6 to estimate power. Recent co-loss was tested in four sister-species pairs (*S. cerevisiae*/*S. kudriavzevii*, *N. castellii*/*N. dairenensis*, *K. africana*/*K. naganishii*, *T. phaffii*/*T. blattae*) by counting partner losses in the same versus the other sister when a gene had been lost in one sister only. Positional coupling replaced partners by the genes at a given distance in the ancestral pillar order, using all POInT pillars, with a null in which genes were placed in random order. Details are in Supplemental Note S10.

### Shared and WGD-specific propensities

Posterior means of each plant gene’s shared liability *L_g_* and WGD-specific liabilities *D_ge_* were computed from the fitted liability-threshold model and the descendant-level data by Gauss–Hermite quadrature (41 nodes for *L_g_*, 21 for each *D_ge_*), integrating each WGD’s specific factor and combining WGDs through *L_g_*. Associations with gene features used genes present in at least two WGDs. Pre-duplication protein-sequence constraint and alignment length came from outgroup alignments that exclude the duplicated lineage; yeast copy-number limits (gTOW) (Makanae et al. 2013) and extra-copy fitness costs (MoBY-CEN) (Morrill and Amon 2019) came from published laboratory experiments. The plant shared propensity of a eukaryotic gene was the mean *L_g_*of its angiosperm lineages; its association with yeast and TGD retention was estimated by partial Spearman correlation (rank residuals) with 1,000 bootstrap resamples.

### Dated tree

Divergence times in Fig. 1A are TimeTree 5 medians with 95% intervals for representative species pairs (Kumar et al. 2022). WGD ages are published estimates: At-*α*, about 47 million years ago (Kagale et al. 2014); grass *ρ*, about 70 (Paterson et al. 2004); legume, about 59 (Schmutz et al. 2010); Maleae, 30–45 (Wu et al. 2013); budding yeast, about 100 (Wolfe and Shields 1997); and TGD, 267–286 (Qi et al. 2024). The stylommatophoran WGD is shared by all sampled species and absent from *Onchidella* (Liu et al. 2021; Chen et al. 2022; McHale et al. 2025), so it predates their common ancestor (about 96 million years ago); its exact age is uncertain. Dates are for orientation and do not enter any analysis (Supplemental Note S8, Supplemental Table S9).

### Statistics and reproducibility

Permutation *P* values were computed as (*b* + 1)/(*B* + 1) (Phipson and Smyth 2010), so a value equal to 1/(*B* + 1) indicates that no permuted statistic reached the observed one. Intervals are two-sided bootstrap percentile intervals unless stated otherwise. Analyses used Python 3 (NumPy, SciPy, pandas, scikit-learn) with fixed random seeds. Every number in this article is written by script from a single values file, and all code, intermediate tables and freeze records are provided (see Software availability).

### Software availability

All analysis scripts, the derived data tables, the source data for every figure and table and the script that writes every number in this article are provided as Supplemental Code. A single command (scripts/run_all.sh) regenerates all results, figures, tables and the manuscript from the frozen reproducibility package, which contains the inputs, source manifests, decision log and freeze records with SHA-256 hashes and is available from Zenodo (https://doi.org/10.5281/zenodo.22896820).

## Data access

No new sequencing data were generated in this study. The genome reconstructions, orthology releases and annotation resources analysed (POInT, OMA, TAIR, BioGRID, PaxDb, Complex Portal, CORUM, CYC2008, TimeTree and the expression atlases cited in Methods) are publicly available from their providers, and the versions and checksums used are recorded in the reproducibility package at Zenodo (https://doi.org/10.5281/zenodo.22896820). The stylommatophoran WGD reconstruction generated in this study is included in the same reproducibility package, and all derived tables and the source data for every figure and table are provided as Supplemental Code.

## Acknowledgments

We thank the developers and maintainers of POInT, OMA, TAIR, BioGRID, PaxDb, TimeTree and the other public genomic and expression resources used in this study. AI-assisted tools were used for code development, analysis implementation and editing under the authors’ direction; all analyses are documented in the reproducibility package for independent checking. This research received no specific grant from any funding agency in the public, commercial or not-for-profit sectors.

## Author contributions

K.S. conceived the study, designed and performed the analyses, curated the data and wrote the manuscript. N.G. contributed to investigation and validation and reviewed and edited the manuscript. Both authors read and approved the final manuscript.

