## Supplementary information for "Duplicate gene retention is partly repeatable across independent whole-genome duplications in plants, fungi and animals"

Contents: Supplemental Notes S1–S10, Supplemental Figures S1–S7 and Supplemental Tables S1–S15. The analysis code and source data are provided separately as Supplemental Code.

### Contents

**About this supplement** **1**

**Note S1. Reconstructions, retention outcomes and orthology** **2**

**Note S2. Robustness to orthology, sampling and dependence** **3**

**Note S3. Liability-threshold model** **3**

**Note S4. Gene-property covariates and adjustment** **4**

**Note S5. Cross-kingdom latent analysis** **5**

**Note S6. Prespecified test protocol and freeze records** **5**

**Note S7. Prespecified tests of individual mechanisms** **7**

**Note S8. Dated tree of the duplications** **7**

**Note S9. Glossary** **7**

**Note S10. Within-genome tests and the components of the shared propensity** **9**

**Supplemental figures** **10**

**Supplemental tables** **14**

### About this supplement

This supplement extends the Methods of the main text with derivations, validation analyses, the complete prespecification record and additional results. Notes S1–S5 give technical detail on reconstruction and orthology (S1), robustness (S2), the liability-threshold model (S3), gene-property features and adjustment (S4) and the cross-kingdom latent analysis (S5). Note S6 describes the staged protocol under which the prespecified tests were run and lists their freeze records (Table S11); Note S7 summarizes the prespecified tests of individual mechanisms; Note S8 documents the dates in Fig. 1A; Note S9 is a glossary; and Note S10 describes the within-genome tests and the decomposition of propensity behind Figs. 6 and 7 of the main text

(Tables S12–S15). Throughout, "lineage" or "ancestral gene" means the single gene of a common ancestor from which a set of modern genes descends, represented by a hierarchical orthologous group (HOG); "event" means one whole-genome duplication (WGD); a pillar is "retained" in a descendant genome when both ohnologs are present; "replay" denotes the association of retention between WGDs; and "coordinate" denotes the taxonomic level (Mesangiospermae, Eukaryota or Bilateria) at which HOGs are defined.

### Note S1. Reconstructions, retention outcomes and orthology

**Unit of analysis.** The unit of inference is an ancestral gene lineage, represented by a hierarchical orthologous group (HOG) at a stated taxonomic level, not a modern gene copy. A lineage contributes to several comparisons; every resampling procedure therefore resamples whole lineages and carries all of their event records together.

**Reconstructions.** POInT pillars represent ancestral loci with two WGD-derived tracks followed through every descendant genome (Hao et al. 2022; Conant 2023). Frozen release files, checksums and parsing audits are part of the reproducibility package. For pillar $i$ in event $e$ the copy count in each descendant is one or two (zero-copy cells are not represented in POInT and were treated as a parsing failure). Retention occupancy $y_{ie} = k_{ie}/m_e$ counts descendants retaining both copies. Occupancy is an observed tip pattern; it does not identify the branch on which a copy was lost. Descendant genomes and releases are listed in Table S2.

**Binary retention.** For the At-$\alpha$, grass $\rho$, legume, yeast and TGD events, a lineage was "retained" when at least one descendant retained both copies. In these old events 27–42% of pillars meet this criterion (Table S10). In the Maleae event, 97% of pillars retain both copies in at least one of five closely related genomes, so this criterion is uninformative; the contrast used was complete retention in all five genomes (27% of pillars). Analyses on the rank scale use all occupancy values and are independent of these thresholds.

**Mesangiospermae mapping.** Translated POInT proteins were matched to OMA All.May2026 proteins by identical normalized peptide sequence within the same species (Altenhoff et al. 2024). For each pillar we recorded the HOGs supported by every exactly matched descendant. A pillar was eligible when at least two descendant species had exact matches, all supported one Mesangiospermae HOG, and no species supported several HOGs. HOG–pillar relations were then required to be one-to-one within an event, and HOGs with within-event multiplicity in any represented reference event were excluded from the primary analyses. None of the Apple/Pear genomes is represented in OMA, so Apple/Pear pillars were mapped with DIAMOND (Buchfink et al. 2021) using a precision-first rule (support from at least two descendant species, unanimous HOG assignment, no within-species conflict, one-to-one relation) calibrated on held-out exact assignments of the reference events before Apple/Pear outcomes were opened.

**Deep and animal coordinates.** TGD and yeast pillars were mapped to OMA Eukaryota HOGs with the same exact, unanimous and one-to-one rules. Angiosperm lineages were lifted to Eukaryota HOGs through the OMA hierarchy; the prespecified plant-score comparisons required a one-to-one lift. The TGD–Stylommatophora comparison used native OMA Bilateria HOGs.

**Stylommatophora reconstruction.** All 12,570 protein-coding *Onchidella celtica* genes on chromosomes 1–18 entered a loss-inclusive candidate universe. Protein similarity nominated up to five candidates per descendant; accepted two-track loci required significant outgroup-chromosome enrichment (at least ten anchors per block), at least two additional compatible anchors within $\pm 20$ genes, an accepted paralogon and cross-descendant block correspondence. High-confidence (618) and moderate-confidence (349) loci were accepted; unphased, multicopy-ambiguous, small-scale duplication, paralogon-conflict and insufficiently supported candidates were excluded. Mapping to OMA was calibrated on a balanced 5,400-protein leave-one-species-out panel from nine molluscan proteomes (precision 0.9994, coverage 0.9957). Two independent phylogenetic checks were made. A post-freeze panel of 100 high-confidence families analysed with outgroup-rooted neighbour-joining trees showed lower within-track than between-track protein distance in 75 families; strict monophyly of both tracks, a demanding criterion for ancient duplicates, was recovered in 25. A separately prespecified panel of 300 loci analysed with maximum-likelihood trees yielded 213 informative trees, of which 8 (3.8%) conflicted with the inferred phase, below the prespecified 20% concern threshold. Because acceptance of a locus requires two-track evidence in at least one descendant, every accepted locus has occupancy $\geq 1/3$, and the structural class is partly determined by retention (all moderate loci have occupancy exactly 1/3). Correlations within the high-confidence subset should therefore be read as a description rather than a test independent of the outcome.

### Note S2. Robustness to orthology, sampling and dependence

**Attrition with taxonomic depth.** At the Mesangiospermae level, 69–79% of reference pillars obtained a one-to-one HOG (Fig. S1a). Populations shrink further in pairwise comparisons because a lineage must be mapped in two events, and at the Eukaryota level because the one-to-one lift fails whenever several angiosperm lineages descend from one eukaryotic ancestral gene. Analysed pillars were retained less often than excluded pillars (for example 36% versus 54% for At-$\alpha$). Excluded pillars comprise those without sufficient exact matches (the largest class), those with within-event multiplicity or HOG conflict, and those whose two retained copies matched different HOGs, all of which are retained by construction (332 pillars for At-$\alpha$); every class is retained more often than analysed pillars, the last by construction (Table S4).

**Event-specific exclusion cannot create association.** Let $R_a, R_b \in \{0, 1\}$ be retention in two events and $S$ inclusion in the analysis. If inclusion depends on each event's outcome separately, $P(S = 1 \mid R_a, R_b) = f(R_a)g(R_b)$, then among included lineages $P(R_a, R_b \mid S = 1) \propto P(R_a, R_b)f(R_a)g(R_b)$, and the cross-product ratio $\frac{P_{11}P_{00}}{P_{10}P_{01}}$ is unchanged because the factors $f$ and $g$ cancel. Exclusion of multiplicity and within-species conflicts is of this form. Exclusion can still change the population to which estimates apply, which is why we also report the relaxed rule.

**Alternative rules.** Table S5 and Fig. S2a report replay under four alternatives. Requiring exact support from at least three species ($T = 0.208$, 1,819 pair observations), averaging the pillars of multiplicity-affected lineages ($T = 0.217$, 3,445), restricting to lineages with a one-to-one Eukaryota projection ($T = 0.182$) and using the Pentapetalae coordinate for At-$\alpha$ and legume ($r = 0.230$) all gave estimates close to the primary $T = 0.210$.

**Descendant sampling.** The three reference events have six, five and four descendants. Recomputing occupancies from every combination of four descendants (15×5×1 = 75 combinations) gave a median global replay of 0.202 (range 0.183–0.216), and pairwise odds ratios between 1.92 and 2.60 (Fig. S2b).

**Dependence.** The three reference correlations share lineages and events. We resampled lineages jointly across all pairs (2,000 resamples), estimated the covariance matrix of the Fisher-transformed pair estimates and pooled them by generalized least squares. The mean correlation among pair estimates was only 0.048, because each pair is dominated by lineages absent from the other pairs. GLS pooling gave 0.210 (95% CI 0.172–0.248) for the reference pairs and 0.190 (0.160–0.220) for all six plant pairs; omitting any one event gave 0.161–0.210 (Fig. S2c).

**Deep populations.** The prespecified deep tests used lineages with a one-to-one projection and at least two reference source events (186 yeast and 151 TGD lineages). Averaging plant scores across all angiosperm lineages of the same Eukaryota HOG enlarged the populations to 277/235 (at least two sources) and 726/567 (at least one source). Yeast AUCs were 0.66 and 0.65; TGD AUCs were 0.59 ($P = 0.019$) and 0.57 ($P = 0.010$).

### Note S3. Liability-threshold model

**Model.** For lineage $g$, WGD $e$ and descendant $d$, $z_{ged} = \tau_{ed} + \lambda_d L_g + \gamma_d D_{ge} + \varepsilon_{ged}$ with $L_g, D_{ge} \sim N(0, 1)$ independent and $\varepsilon_{ged} \sim N(0, 1 - \lambda_d^2 - \gamma_d^2)$; two copies are observed when $z_{ged} > 0$. The thresholds $\tau_{ed}$ absorb differences in overall retention between descendants. $L_g$ is the ancestral lineage's propensity shared by all WGDs, $D_{ge}$ its WGD-specific propensity shared by all descendants of that WGD, and $\varepsilon_{ged}$ everything specific to one descendant (including losses on terminal branches). This is a bifactor model (Holzinger and Swineford 1937) for binary indicators, the standard liability model of quantitative genetics (Falconer 1965).

**Estimation.** Tetrachoric correlations (Pearson 1900) were computed for all 190 pairs of the 20 plant descendants on pairwise-complete lineages with a 0.5 continuity correction. Loadings were obtained by weighted least squares fitting of the 190 off-diagonal correlations (weights: shared lineages), constrained to $[0, 1)$. Per-descendant loadings are shown in Fig. S3. Event summaries average $\lambda_d^2$ and $\gamma_d^2$ over the descendants of each event. Repeatability $\bar{\lambda}^2/(\bar{\lambda}^2 + \bar{\gamma}^2)$ (Nakagawa and Schielzeth 2010) is the expected correlation between a lineage's WGD-level propensities (shared plus WGD-specific components) in two independent WGDs.

**Interpretation of the components.** The event-specific component is large partly because descendants of one WGD share most of their post-duplication history; for At-$\alpha$, whose descendants are closely related Brassicaceae, within-event tetrachoric correlations average 0.86. Phylogenetic structure among descendants within an event is absorbed into $D_{ge}$ and $\varepsilon_{ged}$ and does not affect the shared component, which is identified only from correlations between descendants of different WGDs. If close relatives inflate the within-event correlations, the repeatability is underestimated, so the reported value is conservative.

**Disattenuation and ceiling.** The latent correlation between events $a$ and $b$ is the mean cross-event tetrachoric correlation divided by the geometric mean of the mean within-event correlations (Spearman 1904). The ceiling simulation keeps the fitted thresholds and the total gene-level variance ($\lambda_d^2 + \gamma_d^2$) of every descendant but assigns all of it to the shared factor; the resulting correlation is the largest that the observed incidence, number of descendants and retention frequencies allow if lineage propensity were identical across WGDs.

**Uncertainty.** Lineages were resampled 1,000 times with replacement; for each resample the tetrachoric matrix was recomputed and the model refitted. Intervals are 2.5th and 97.5th percentiles.

**Parameter recovery.** We simulated data with the same structure (four events with six, five, four and five descendants, 4,000 lineages with event-specific incidence of 30–55%, descendant-specific thresholds) under known components and refitted the model (20 replicates per scenario; script `14_latent_simulation_check.py`). True repeatabilities of 0.17, 0.31 and 0.43 were recovered as 0.19, 0.31 and 0.43 on average (replicate s.d. 0.02–0.03), and the shared and event-specific shares were recovered to within 0.015.

### Note S4. Gene-property covariates and adjustment

**Sources.** GO, GO-slim, mutant phenotypes, gene aliases and publication links: TAIR public data release of 31 March 2018 (files `ATH_GO_GOSLIM.txt`, `Locus_Germplasm_Phenotype_20180330.txt`, `gene_aliases_20180330.txt`, `Locus_Published_20180330.txt`) (Berardini et al. 2015); SHA-256 checksums of the files used are recorded in the package. Physical interactions: BioGRID 5.0.261 multivalidated physical interactions (Oughtred et al. 2021). Protein abundance: PaxDb integrated whole-organism data set for *A. thaliana* (Wang et al. 2015). Aneuploidy and ploidy expression responses: ERCC-normalized RPKM matrix of GSE79676 (Hou et al. 2018). Complex membership: Complex Portal, CORUM and CYC2008 mapped to Eukaryota HOGs, with conserved membership requiring qualifying evidence in more than one major eukaryotic lineage (Meldal et al. 2022; Giurgiu et al. 2019; Pu et al. 2009). Nonfocal expression atlases: *Amborella* TPM, grapevine array and cacao CPM data summarized within species (Flores-Tornero et al. 2020; Fasoli et al. 2012; Kulesza et al. 2024). Nonfocal sequence constraint: Poisson-corrected protein distances among single-copy orthologs of *Amborella*, grapevine, cacao and peach, expressed relative to genome-wide medians.

**Linking to lineages.** *A. thaliana* genes were linked to Mesangiospermae HOGs through exact OMA matches of At-$\alpha$ pillar genes (5,151 of 8,631 lineages); Entrez and UniProt cross-references from OMA linked genes to BioGRID and PaxDb. Eukaryota-level features were attached through the OMA lift. Essentiality was assigned when a TAIR phenotype description matched lethal, embryo-defective, embryo-arrest, seed-abortion or gametophyte-lethal terms, or when the gene carried a SeedGenes *EMB* symbol (Meinke et al. 2008; Lloyd and Meinke 2012).

**Design.** Features with missing values were median-imputed and accompanied by a missingness indicator; all columns were standardized. The 76 columns comprise 45 GO-slim indicators and experimental GOterm count (function); three family-size and duplication-history measures; phylostratum, sequence constraint and alignment length (conservation and age, with indicators); interaction degree and publication count; two essentiality indicators; five dosage measures; and four expression and abundance measures, with indicators.

**Adjustment variants.** *In-sample* ordinary least squares removes everything linearly predictable from the features, including chance alignment; its covariance decomposition is exact, and we subtract the mean fraction obtained after permuting feature rows (200 permutations), which estimates the chance contribution (for example 5–8% for the large At-$\alpha$ pairs). *Cross-fitted* residuals are obtained from models fitted to other folds and remove only generalizable covariate effects (Chernozhukov et al. 2018). The *ridge* variant adds 488 GO terms with the penalty chosen by internal cross-validation (Hoerl and Kennard 1970). Because measured covariates are imperfect proxies, all variants under-adjust for the underlying properties, and the residual replay is an upper bound on the component independent of those properties.

**Adjusted odds ratios.** With 76 features and a few hundred lineages per pair, direct logistic adjustment is unstable. For each pair we therefore estimated, by five-fold cross-fitting, a ridge-penalized logistic risk score for retention in each event from the features, and fitted a logistic regression of retention in one event on retention in the other plus both risk scores.

**Within-family replay.** Lineages were grouped by OMA root family; for each pair, families with at least two lineages present in both events were retained. Each event's percentiles were centred on family means, the centred values correlated, and significance assessed by permuting the second event's centred values within families (5,000 permutations). The pooled estimate combines the three reference pairs with $(n-3)$ weights.

**Predictive comparison.** See Table S7. The lineage score involves no fitted parameters and is therefore not advantaged by overfitting; the covariate and combined models are cross-fitted.

### Note S5. Cross-kingdom latent analysis

At the Eukaryota coordinate, each plant event's lineages were lifted one-to-one and combined with the descendant-level copy states of the TGD (8 genomes) and yeast WGD (11 genomes), giving 39 descendant genomes. Tetrachoric correlations and disattenuated pairwise latent correlations were computed as in Note S3 (Fig. S7; Table S8). Latent correlations between the three reference plant WGDs and the yeast WGD were 0.37, 0.32 and 0.41, and between the TGD and yeast WGD 0.31 (95% CI 0.08–0.45). These values are comparable to the latent correlations among angiosperm WGDs at the Mesangiospermae coordinate, indicating that the shared propensity is not confined to plants. Lineages that can be followed from angiosperms to fungi and animals are older and more conserved than average, so these estimates apply to that subset. A hierarchical extension with separate pan-eukaryotic and plant-specific factors was explored but is weakly identified with only two non-plant events and is not interpreted.

### Note S6. Prespecified test protocol and freeze records

**Organization.** The study was run as a sequence of milestones recorded in a decision log (`DECISION_LOG.md` in the reproducibility package). Each milestone had a written authorization that fixed, before any outcome it would use was opened, the input files and their SHA-256 hashes, the analysis population, the statistic, the numbers of permutations and bootstrap resamples, the random seeds and the decision gates. Results were labelled by applying the gates without discretion, and a failed gate could not be rescued by a secondary or sensitivity analysis. The log and all freeze artefacts were kept in a version-controlled repository whose commits time-stamp each freeze. The protocol was internal; the study was not registered with an external registry. Table S11 lists the freeze records of the tests reported in Table 2 of the main text.

**Reference replay (milestone M4).** Retention outcomes (milestone M2) and the exact Mesangiospermae map (M3B) were frozen independently and joined for the first time in M4, after their hashes, seeds and gates had been recorded. Gate 1 required a positive $T_{\text{replay}}$, one-sided permutation $P \leq 0.001$ and a bootstrap lower bound above zero; Gate 2 required $T_{\text{replay}} \geq 0.10$; Gate 3 required positive leave-one-WGD-out transfer with

$P \leq 0.01$ on the 337 HOGs present in all three reference events. All three gates were met ($T_{\text{replay}} = 0.210$, 0.172–0.248; $T_{\text{LOO}} = 0.221$; both $P = 1/100{,}001$).

**Apple–pear (M5A–M5B).** M5A calibrated the DIAMOND mapping of Maleae proteins on held-out exact assignments of the reference events, without access to any Maleae outcome, and froze the map. M5B intersected the map with the reference universe, computed the unweighted mean of the available reference percentiles (at least two required) and froze and committed the prediction table before the Maleae outcome was opened. Gate V1 required $\rho > 0$, one-sided $P \leq 0.01$ and a bootstrap lower bound above zero; Gate V2 required $\rho \geq 0.10$. Both were met ($\rho = 0.169$, 0.069–0.262; $P = 0.00047$; $n = 373$).

**Distant targets (M6A–M6E).** The order of the distant targets was fixed when M6A, an audit of the feasibility of exact Eukaryota-level mapping, was authorized: the TGD was evaluated first (M6B–M6C) and the yeast WGD second (M6D–M6E). For each target, Stage 1 froze and committed the plant-derived prediction, requiring a one-to-one Eukaryota bridge and at least two reference ranks; only then did Stage 2 construct the target outcome from both POInT representations, require their exact agreement and rank occupancy over the complete exact-supported deep universe of the target (1,187 TGD and 1,392 yeast HOGs). Gates C1 (TGD) and Y1 (yeast) required $\rho > 0$, one-sided $P \leq 0.05$ and a bootstrap lower bound above zero; Gates C2 and Y2 additionally required $\rho \geq 0.10$ and $P \leq 0.01$. The TGD test failed C1 ($\rho = 0.107$, −0.050 to 0.260; $P = 0.094$; $n = 151$) and was recorded as a negative result. The yeast predictor and rules were not changed after this result, and the yeast test met Y1 and Y2 ($\rho = 0.280$, 0.131–0.418; $P = 0.00009$; $n = 186$). Neither the apple–pear nor the TGD outcome entered any later predictor.

**Animal comparisons (M11A–M11B-R, M13A–M13B).** The TGD–Stylommatophora comparison was specified after the plant-to-target tests and went through four versions on 3 September 2026, each frozen before its own step. M11A (Metazoa HOGs; at least 200 shared one-to-one units; $\rho \geq 0.10$) and M11A-R (Bilateria HOGs; at least 200 units) stopped at 170 and 160 units in identifier-only audits, before any snail retention value had been computed. M11B (at least 150 complete pairs; strong replay $r \geq 0.25$) stopped at 146 complete pairs before any association was calculated. M11B-R then froze the 146 HOGs, rank hashes and gates, with no sample-size gate, before computing the correlation once; strong replay required $r \geq 0.25$, one-sided $P \leq 0.01$ and a bootstrap lower bound above zero. The association was positive and met the inferential criteria but fell below the effect threshold ($r = 0.226$, 0.061–0.385; $P = 0.0033$). The strict comparison of the plant score with snail retention (M13A–M13B) froze a universe of 25 Eukaryota HOGs; at this size the correlation detectable with 80% power was about 0.49 at one-sided $\alpha = 0.05$, and the result was unresolved ($r = 0.033$, −0.303 to 0.340; $P = 0.44$).

**Effect-size thresholds declared at the outset.** The protocol of 27 August 2026 also declared effect-size thresholds for leave-one-WGD-out prediction among the four plant WGDs measured by AUC: fail below a mean of 0.60 (or two events at or below 0.55), weak from 0.60 to below 0.65, pass at a mean of at least 0.65 with every event above 0.58, and strong pass at a mean of at least 0.70. The executed M4 design replaced these with the rank-correlation gates above, and AUC was not computed at the time. Applied after the fact to the leave-one-WGD-out AUCs of the history score reported in the main text (0.58–0.63; mean 0.61; Table S7), these thresholds would classify within-plant prediction as weak. The same protocol prohibited describing evolution as predictable on the basis of the four plant WGDs alone; the manuscript therefore describes retention as partly repeatable and the history score as carrying transferable information.

**Analyses that were not prespecified.** The following analyses—conditional retention probabilities, risk and odds ratios, the liability-threshold decomposition, covariate adjustment, the predictive comparison, many-to-one aggregation for the deep targets, rarefaction, generalized least squares pooling, alternative orthology maps and the within-genome tests of Note S10—were designed after the prespecified results were known. They use the same frozen outcomes and maps and are reported as estimation and sensitivity analyses rather than confirmatory tests.

### Note S7. Prespecified tests of individual mechanisms

After the target tests (Note S6) and before the estimation analyses listed in Note S6, we tested individual mechanistic proxies under the same staged protocol (milestones M7 onward). The hypotheses were formulated after the replay and transfer results were known (the decision log describes this phase as post-result mechanism discovery), but each test's features, populations, orientation and decision rules were fixed before its own outcome was examined, and failures were retained (records and hashes in the package). Results are summarized in Fig. S6. The yeast fitness cost of a single extra gene copy (MoBY-CEN, 429 lineages) (Morrill and Amon 2019) was not associated with retention ($\rho = 0.050$, 95% CI $-0.048$ to $0.148$). The median absolute transcriptional response of *Arabidopsis* genes to trisomies of other chromosomes (Hou et al. 2018) was not associated with retention in the reference WGDs ($\rho = -0.013$, $-0.090$ to $0.065$). Nonfocal protein-sequence constraint was not associated ($\rho = 0.036$, $-0.058$ to $0.130$). Conserved expression magnitude was positively associated ($\rho = 0.125$, $0.042$ to $0.206$; Holm-adjusted $P = 0.0063$), whereas expression breadth was not ($\rho = 0.041$, $-0.054$ to $0.137$). Adjusting replay for expression magnitude on identical lineage sets changed the Fisher-scale replay by $\Delta Z = 0.0068$ ($-0.0168$ to $0.0327$). A four-property model fitted on two reference WGDs and evaluated on the third did not outperform single properties ($\rho = 0.048$, $-0.146$ to $0.162$; 185 lineages). An exploratory test of whether conserved complex membership strengthened transfer to TGD and yeast gave an interaction of 0.050 ($-0.207$ to $0.322$). The broader adjustment in the main text is consistent with these results and extends them to many more properties and lineages.

### Note S8. Dated tree of the duplications

The tree in Fig. 1A of the main text shows the established relationships among the lineages studied: plants and opisthokonts (fungi plus animals) diverged first; fungi then separated from animals; and, among animals, deuterostomes (including fishes) separated from protostomes (including snails). Among flowering plants, monocots (grasses) separated from eudicots, and among eudicots the mustard family (malvid rosids) separated from the fabid rosids, which include the legumes and the rose family (apple and pear). Node ages are TimeTree 5 medians for representative species pairs, with 95% intervals (Kumar et al. 2022); they were retrieved in October 2026 and are listed with the species used in Table S9. Each coloured wedge spans the crown age of the sampled descendant genomes, that is, the time since they last shared an ancestor.

Duplication ages are published estimates (Table S9): the At-$\alpha$ duplication about 47 million years ago (Kagale et al. 2014), the grass $\rho$ duplication about 70 (Paterson et al. 2004), the papilionoid legume duplication about 59 (Schmutz et al. 2010), the apple–pear duplication 30–45 (Wu et al. 2013), the budding-yeast duplication about 100 (Wolfe and Shields 1997) and the teleost duplication 267–286 (Qi et al. 2024); earlier estimates for the teleost duplication were older (about 320–350 million years), and the figure uses the most recent dated analysis. Every duplication age is older than the crown age of the corresponding sampled genomes, as expected for a duplication shared by all of them. For the snail duplication, duplicated Hox clusters and macrosynteny are shared by all sampled stylommatophorans but absent from *Onchidella* (Liu et al. 2021; Chen et al. 2022; McHale et al. 2025), so the duplication is older than the sampled crown (about 96 million years). A synonymous-divergence estimate of about 70 million years (Liu et al. 2021) is younger than this crown and was not used; the figure therefore marks the snail duplication with an open star and indicates only that it is older than 96 million years. The time axis is logarithmic, $\log_{10}(t + 10)$ with $t$ in millions of years, so that divergences more than a billion years old and duplications a few tens of millions of years old are both legible. Dates are shown for orientation only and do not enter any analysis.

### Note S9. Glossary

**Whole-genome duplication (WGD), polyploidy.** Doubling of an entire genome in one step, by genome doubling within a species (autopolyploidy) or by hybridization between species (allopolyploidy).

**Ohnologs.** The pair of gene copies produced by a WGD.

**Fractionation; biased fractionation.** The loss of one ohnolog from most pairs after a WGD; biased fractionation is the preferential retention of genes from one parental subgenome after allopolyploidy.

**Retained / returned to single copy.** In a descendant genome, an ancestral locus is retained when both ohnologs are present. Here a locus is retained after a WGD when at least one descendant genome retains it (apple–pear: all five genomes).

**Retention occupancy.** The fraction of a WGD's descendant genomes that retain both ohnologs of a locus.

**Event-wide percentile.** The rank of a locus's retention occupancy among all mapped loci of the same WGD, computed once and not recomputed within pairwise intersections.

**Ancestral gene, lineage, HOG.** The single gene of a common ancestor from which a set of modern genes descends. Hierarchical orthologous groups (HOGs) in the OMA database collect these genes for a named ancestor (for example Mesangiospermae, Eukaryota or Bilateria).

**Pillar.** In POInT reconstructions, one ancestral locus followed through all descendant genomes of a WGD as two duplicate tracks.

**One-to-one mapping.** Accepting a match between a pillar and a HOG only when neither is matched to anything else within the event.

**Reference and target WGDs.** The three angiosperm WGDs used to build predictions (At-$\alpha$, grass $\rho$, legume), and the WGDs on which those predictions were tested (apple–pear, budding yeast, TGD).

$T_{\text{replay}}$**.** The $(n - 3)$-weighted Fisher mean of pairwise correlations of event-wide percentiles among the reference WGDs.

**Risk ratio, odds ratio.** The risk ratio compares the probability of retention after one WGD among loci retained versus returned to single copy after another; the odds ratio compares the corresponding odds. Values above 1 mean that retention tends to repeat.

**AUC.** Area under the receiver operating characteristic curve: the probability that a randomly chosen retained locus receives a higher score than a randomly chosen locus returned to single copy (0.5 = chance, 1 = perfect).

**Tetrachoric correlation.** The correlation between two binary outcomes estimated on the scale of an underlying continuous liability.

**Liability-threshold (bifactor) model.** A model in which an unobserved continuous liability determines a binary outcome when it exceeds a threshold; here the liability is divided into components shared across WGDs, specific to one WGD and specific to one genome.

**Repeatability.** The shared component as a fraction of gene-level (shared plus WGD-specific) liability variance; the expected correlation of a gene's propensity between two independent WGDs.

**Disattenuated (latent) correlation.** A between-WGD correlation divided by the geometric mean of the within-WGD correlations, which corrects for the sampling of descendants.

**Attainable maximum.** The largest correlation between two WGDs that the observed sampling and retention rates allow if all gene-level variation were shared, obtained by simulation from the fitted model.

**Cross-fitting.** Estimating a model on part of the data and applying it to the remaining part, so that adjustment removes only effects that generalize.

**Within-genome (fixed-effects) design.** A comparison of the same ancestral gene across the genomes descended from one WGD, with gene and genome fixed effects, so that no property of the gene can account for an association.

**Coupling coefficient $\theta$.** In the within-genome design, the change in the probability that a genome retains both copies of a gene per unit change in the fraction of its partners (or neighbours) retained in that genome.

**Prespecified test; freeze record; gate.** A test whose population, prediction, statistic and decision rules (gates) were fixed before the outcome was examined; the freeze record is the hashed and committed file that documents this.

### Note S10. Within-genome tests and the components of the shared propensity

**Rationale.** Associations between gene properties and retention cannot separate causes from correlates, because any measured property may stand in for an unmeasured one. Within one WGD, however, the same ancestral gene is retained in some descendant genomes and lost in others. Comparing these genomes holds the gene constant, so that its function, expression, family size, constraint and any unmeasured property are fixed by design. The question becomes whether the state of something else in the same genome, such as the gene's complex partners or its chromosomal neighbours, predicts whether that genome lost the gene.

**Model.** For gene $g$ and genome $d$ of one WGD, $y_{gd} \in \{0, 1\}$ records retention of both ohnologs, and $P_{gd}$ is the fraction of the gene's partners (or neighbours) retained in $d$. We fitted $y_{gd} = \alpha_g + \beta_d + \theta P_{gd} + e_{gd}$ by least squares after two-way demeaning of the balanced gene-by-genome panel. Genes whose state does not vary contribute through the genome means, as in any fixed-effects estimator. Because $y$ is binary and the panel is short, the estimator's sampling distribution is not assumed: inference uses permutation nulls that keep each gene's data and break only the partner relationships, and bootstrap intervals over focal genes. With few genomes per WGD the fixed-effects estimator has a small negative bias, which the permutation null shares; $z$ is therefore always computed against the null mean.

**Partners.** Stable heteromeric complexes from CYC2008, CORUM and Complex Portal (complex_evidence table of milestone M7B) were mapped to Eukaryota HOGs. Two genes of one WGD were partners when they shared at least one complex, unless they were paralogous lineages of one eukaryotic gene or lay within 20 pillars of each other in the ancestral order. For plant WGDs, Gene Ontology cellular-component terms for stoichiometric complexes (cytosolic ribosome, proteasome, mediator, complex I, spliceosome, snoRNP, small-subunit processome, heterotrimeric G protein, SNARE) gave a second, independent partner definition.

**Results.** Table S12 gives the test for every WGD, Table S13 the yeast robustness, specificity, class, timing, dose-moderation and power analyses, and Table S14 the positional analyses. The yeast coupling is robust to every variation tried; it is concentrated in, but not confined to, the cytosolic ribosome; and it is specific to physical partners. The absence of coupling in the plant WGDs and the TGD is not a matter of power: subsamples of the yeast data with as few genomes as the plant WGDs detect the yeast effect in nearly all cases, and every plant and fish interval excludes the yeast estimate.

**Interpretation and limits.** The design excludes confounding by gene properties but not by causes that act on a whole complex within a lineage at once (for example, selection on growth rate acting on all ribosomal proteins). The sister-pair analysis shows that coupled losses occur on the same recent branches; it could not establish that partner loss precedes gene loss. Ribosomal-protein ohnologs are kept nearly identical by gene conversion, so collapsed assemblies could in principle mimic coupled losses; the result is robust to omitting each genome and is specific to physical partners rather than to near-identical gene families in general. A direct test is available: dosage balance predicts alleviating genetic interactions between deletions of single duplicate copies of partner genes, which can be examined in yeast genetic-interaction maps (Costanzo et al. 2016; Kuzmin et al. 2020).

**Shared and WGD-specific propensities.** Posterior means were computed under the fitted bifactor liability model (Note S3) for each plant gene, integrating each WGD-specific factor by 21-node and the shared factor by 41-node Gauss–Hermite quadrature. For genes observed in only one WGD the two components cannot be separated, so feature associations use genes present in at least two WGDs (Table S15). Pre-duplication constraint and length come from the outgroup alignment panels of milestone M18A (YEAST_PRE_WGD, TGD_PRE_EVENT, PLANT_COMMON), which exclude the duplicated lineage; gTOW and MoBY-CEN values come from the published experiments as frozen in the mechanism programme.

### Supplemental figures

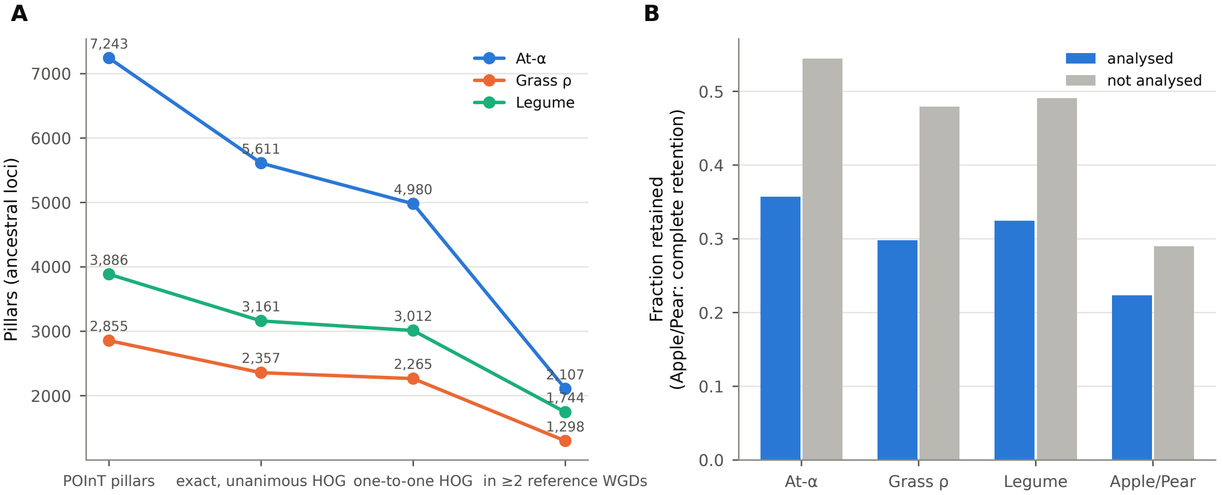


**Figure S1. Orthology attrition and representativeness. A** Pillars of each reference WGD at successive mapping steps at the Mesangiospermae coordinate. **B** Fraction of pillars retained among analysed and excluded pillars (Maleae: complete retention).

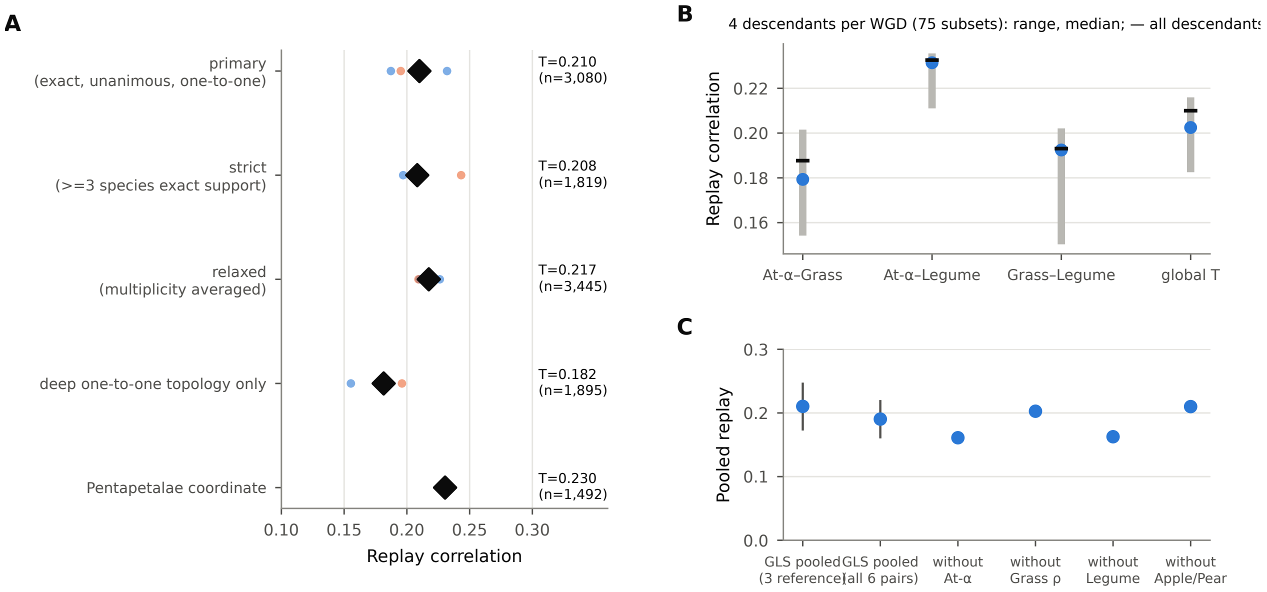


**Figure S2. Robustness of replay. A** Pairwise (small dots, coloured by the first event of the pair) and pooled (diamonds) replay under alternative orthology rules. **B** Replay after rarefying each reference WGD to four descendant genomes (75 combinations): grey bars show the range, blue dots the median and black ticks the estimate with all descendants. **C** Generalized least-squares pooled replay accounting for dependence among pairwise estimates (95% CI), and Fisher-pooled replay omitting each WGD in turn.

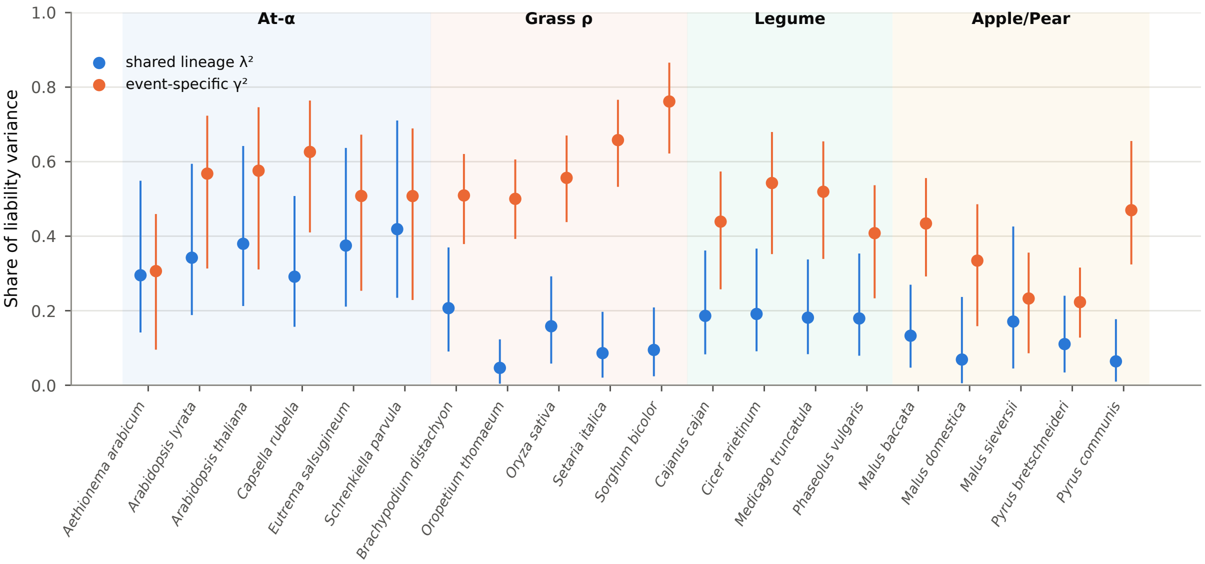


**Figure S3. Descendant-level estimates of the liability model.** Shared lineage ($\lambda_d^2$, blue) and event-specific ($\gamma_d^2$, orange) shares of liability variance for each of the 20 descendant genomes, with 95% bootstrap intervals (1,000 lineage resamples).

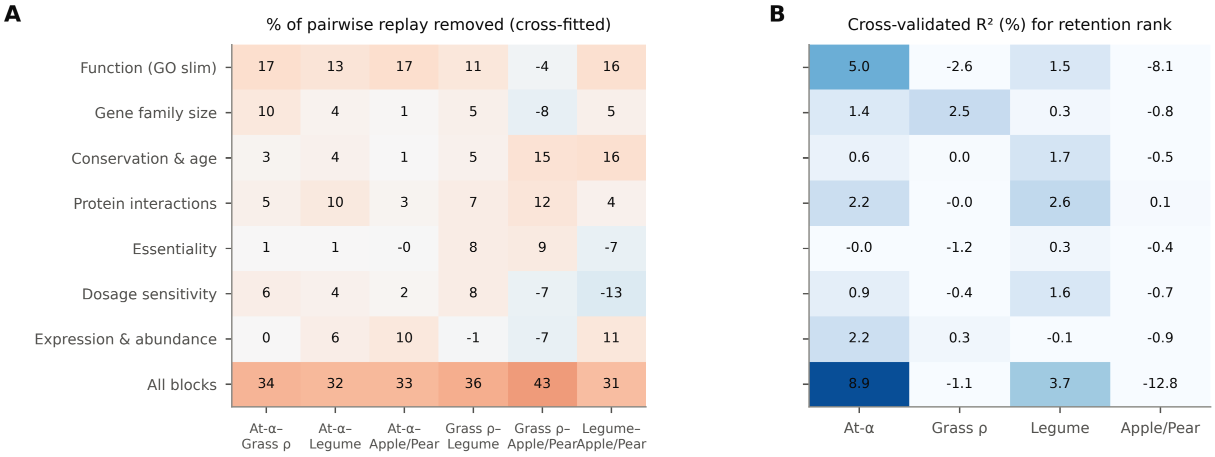


**Figure S4. Covariate blocks. A** Percentage of replay removed by cross-fitted adjustment for each block, for each pair of plant WGDs (negative values: adjustment increased the estimate). **B** Cross-validated $R^2$ of each block for retention rank in each WGD.

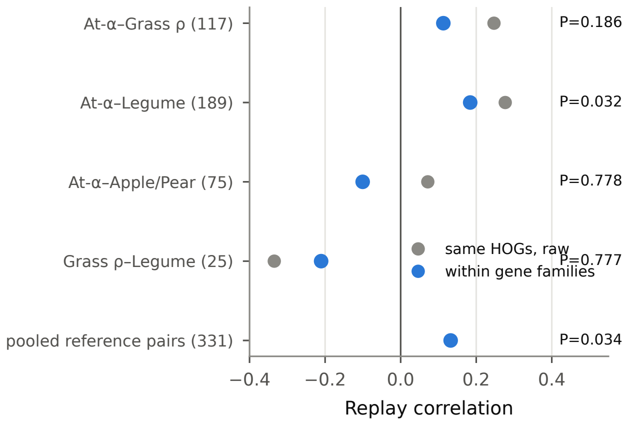


**Figure S5. Replay within gene families.** Replay among lineages of the same OMA root family after centring on family means (blue), compared with the uncentred correlation on the same lineages (grey). $P$: within-family permutation test. Numbers of lineages in parentheses.

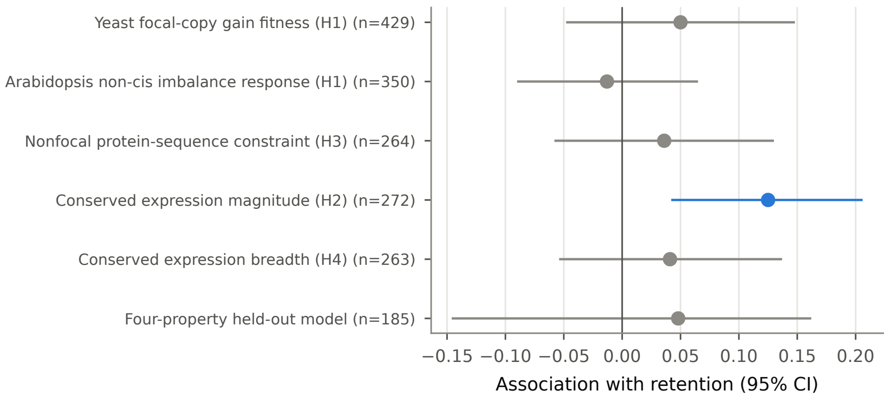


**Figure S6. Prespecified single-mechanism tests** (Note S7). Associations with 95% bootstrap intervals; blue, supported under the prespecified criterion.

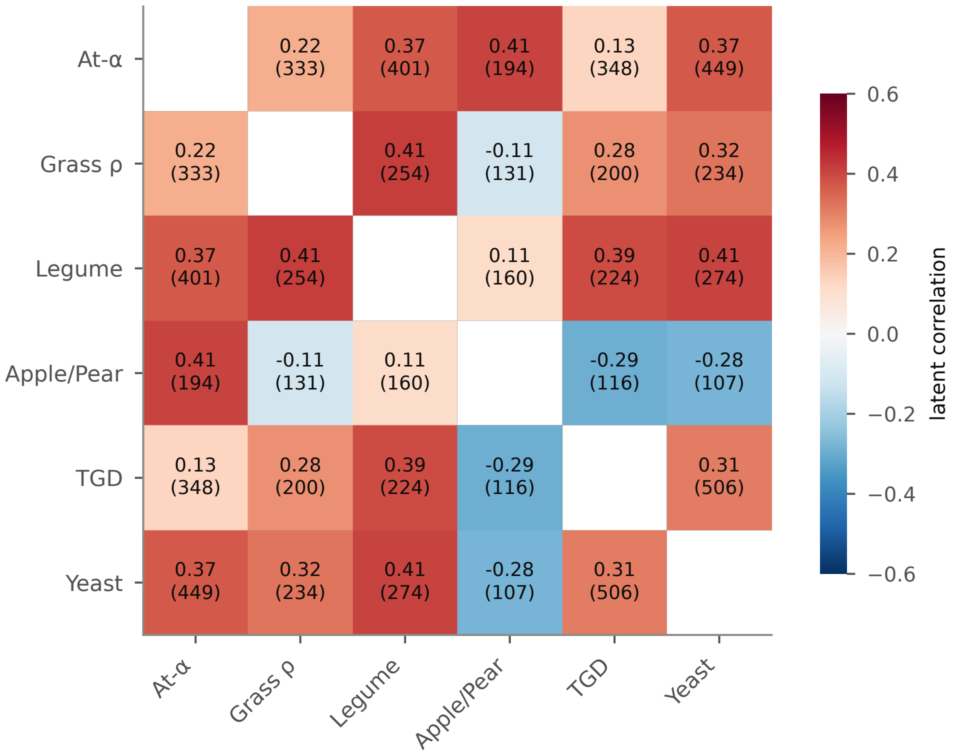


**Figure S7. Cross-kingdom latent correlations at the Eukaryota coordinate.** Pairwise latent correlations of retention propensity (cross-event tetrachoric correlation disattenuated for within-event agreement) among four plant WGDs, the TGD and the yeast WGD; cells show estimates and numbers of shared lineages.

### Supplemental tables

**Table S1. Conditional retention between pairs of plant WGDs.** Cells: retained in both / A only / B only / neither. $P(B \mid A)$ and $P(B \mid \bar{A})$: probability of retention after WGD B given retention or loss after WGD A. AUC: retention occupancy in A discriminating retained from lost lineages in B. 95% intervals from 10,000 lineage bootstraps.

| Pair | $n$ | Cells | $P(B \mid A)$ | $P(B \mid \bar{A})$ | Risk ratio | Odds ratio | AUC | Spearman $\rho$ |
|---|---|---|---|---|---|---|---|---|
| At-$\alpha$ → Grass $\rho$ | 999 | 156/209/162/472 | 0.43 | 0.26 | 1.67 (1.40–2.01) | 2.17 (1.66–2.88) | 0.60 (0.56–0.63) | 0.188 (0.124–0.252) |
| At-$\alpha$ → Legume | 1,445 | 274/277/257/637 | 0.50 | 0.29 | 1.73 (1.52–1.98) | 2.45 (1.97–3.07) | 0.62 (0.59–0.64) | 0.233 (0.180–0.284) |
| At-$\alpha$ → Maleae (Apple/Pear) | 813 | 75/173/126/439 | 0.30 | 0.22 | 1.36 (1.05–1.72) | 1.51 (1.07–2.11) | 0.54 (0.50–0.58) | 0.165 (0.101–0.228) |
| Grass $\rho$ → Legume | 636 | 99/105/121/311 | 0.49 | 0.28 | 1.73 (1.41–2.14) | 2.42 (1.72–3.46) | 0.60 (0.56–0.64) | 0.194 (0.114–0.273) |
| Grass $\rho$ → Maleae (Apple/Pear) | 315 | 27/52/68/168 | 0.34 | 0.29 | 1.19 (0.78–1.68) | 1.28 (0.71–2.19) | 0.53 (0.47–0.59) | 0.066 (-0.047–0.177) |
| Legume → Maleae (Apple/Pear) | 532 | 63/104/83/282 | 0.38 | 0.23 | 1.66 (1.26–2.17) | 2.06 (1.38–3.07) | 0.58 (0.53–0.62) | 0.172 (0.094–0.249) |

**Table S2. Descendant genomes and reconstruction releases.**

| WGD | Release | Descendant genomes |
|---|---|---|
| At-$\alpha$ | AtAlpha-6 | *Aethionema arabicum*, *Arabidopsis lyrata*, *A. thaliana*, *Capsella rubella*, *Eutrema salsugineum*, *Schrenkiella parvula* |
| Grass $\rho$ | GrassRho-5 | *Brachypodium distachyon*, *Oropetium thomaeum*, *Oryza sativa*, *Setaria italica*, *Sorghum bicolor* |
| Legume | LegumeWGD-4 | *Cajanus cajan*, *Cicer arietinum*, *Medicago truncatula*, *Phaseolus vulgaris* |
| Maleae (Apple/Pear) | ApplePearWGD-5 | *Malus baccata*, *M. domestica*, *M. sieversii*, *Pyrus bretschneideri*, *P. communis* |
| Yeast | YeastWGD-11 | *Candida glabrata*, *Kazachstania africana*, *K. naganishii*, *Naumovozyma castellii*, *N. dairenensis*, *Saccharomyces cerevisiae*, *S. kudriavzevii*, *S. uvarum*, *Tetrapisispora blattae*, *T. phaffii*, *Vanderwaltozyma polyspora* |
| TGD | TGD-8 | *Astyanax mexicanus*, *Danio rerio*, *Gasterosteus aculeatus*, *Oreochromis niloticus*, *Oryzias latipes*, *Takifugu rubripes*, *Tetraodon nigroviridis*, *Xiphophorus maculatus* |
| Stylommatophora | this study | *Arion vulgaris*, *Lissachatina fulica*, *Bradybaena similaris* (pre-duplication anchor *Onchidella celtica*) |

**Table S3. Marginal association of selected gene properties with retention rank.** Spearman $\rho$ between each property and the event-wide retention percentile, among lineages with *A. thaliana* members ($^{*}P < 0.01$; $^{**}P < 0.001$; two-sided).

| Block | Property | At-$\alpha$ | Grass $\rho$ | Legume | Maleae |
|---|---|---|---|---|---|
| Function | GO slim: transcription factor activity | +0.070** | +0.064 | +0.107** | +0.046 |
| Function | GO slim: kinase activity | +0.036 | +0.038 | +0.066* | -0.005 |
| Function | GO slim: signal transduction | +0.061** | +0.052 | +0.127** | -0.001 |
| Function | GO slim: protein binding | +0.061** | +0.027 | +0.105** | +0.045 |
| Function | GO slim: ribosome | +0.040* | +0.044 | -0.004 | -0.027 |
| Function | GO slim: chloroplast | -0.043* | -0.035 | -0.132** | -0.055 |
| Gene family size | angiosperm lineages in OMA root family | +0.060** | +0.056 | +0.056 | +0.014 |
| Gene family size | other *A. thaliana* genes in family | +0.058** | +0.069 | +0.064 | +0.022 |
| Gene family size | nested duplication nodes in HOG identifier | +0.104** | +0.164** | +0.064 | +0.065 |
| Conservation & age | phylostratum (0 = oldest) | +0.025 | -0.004 | +0.100** | -0.005 |
| Conservation & age | nonfocal protein-sequence constraint | +0.140* | +0.016 | -0.014 | +0.132 |
| Protein interactions | BioGRID physical degree | +0.107** | +0.087* | +0.072* | +0.062 |
| Protein interactions | TAIR publications (log) | +0.152** | +0.084* | +0.167** | +0.044 |
| Essentiality | lethal / embryo-defective phenotype | -0.032 | -0.024 | -0.039 | -0.027 |
| Essentiality | any recorded mutant phenotype | +0.001 | +0.009 | +0.053 | +0.003 |
| Dosage sensitivity | non-cis aneuploidy expression response | +0.081** | +0.015 | +0.153** | -0.067 |
| Dosage sensitivity | cis deviation from proportional response | +0.126** | +0.104** | +0.128** | -0.010 |
| Dosage sensitivity | conserved heteromeric complex member | +0.086** | +0.079 | +0.121* | -0.009 |
| Expression & abundance | *A. thaliana* seedling expression | -0.007 | +0.021 | -0.039 | +0.016 |
| Expression & abundance | PaxDb protein abundance | -0.005 | +0.048 | -0.062 | +0.001 |
| Expression & abundance | conserved atlas expression magnitude | +0.244** | +0.029 | +0.158 | +0.256 |
| Expression & abundance | conserved atlas expression breadth | +0.159** | -0.039 | -0.008 | +0.309 |

**Table S4. Mapping status and retention of reconstructed pillars.** Analysed: pillars entering the one-to-one Mesangiospermae map used here. Retention is lower among analysed than excluded pillars: pillars whose two retained copies matched different HOGs ("multiple HOGs within species") are retained by construction, and unmapped pillars and those with within-event multiplicity are also retained more often. Exclusions that depend on each event's outcome separately do not alter cross-event odds ratios (Note S2).

| WGD | Set or mapping status | Pillars | Mean occupancy | Fraction retained |
|---|---|---|---|---|
| At-$\alpha$ | analysed | 4,934 | 0.229 | 0.357 |
| At-$\alpha$ | not analysed | 2,309 | 0.361 | 0.544 |
| At-$\alpha$ | excluded within event multiplicity | 631 | 0.298 | 0.444 |
| At-$\alpha$ | hog conflict | 82 | 0.159 | 0.366 |
| At-$\alpha$ | insufficient exact support | 74 | 0.394 | 0.716 |
| At-$\alpha$ | multiple hogs within species | 332 | 0.695 | 1.000 |
| At-$\alpha$ | primary exact unanimous | 4,980 | 0.230 | 0.359 |
| At-$\alpha$ | unmapped | 1,144 | 0.310 | 0.469 |
| Grass $\rho$ | analysed | 2,158 | 0.141 | 0.298 |
| Grass $\rho$ | not analysed | 697 | 0.230 | 0.479 |
| Grass $\rho$ | excluded within event multiplicity | 92 | 0.196 | 0.424 |
| Grass $\rho$ | hog conflict | 24 | 0.142 | 0.333 |
| Grass $\rho$ | insufficient exact support | 99 | 0.135 | 0.384 |
| Grass $\rho$ | multiple hogs within species | 21 | 0.524 | 0.952 |
| Grass $\rho$ | primary exact unanimous | 2,265 | 0.148 | 0.311 |
| Grass $\rho$ | unmapped | 354 | 0.233 | 0.472 |
| Legume | analysed | 2,891 | 0.161 | 0.324 |
| Legume | not analysed | 995 | 0.248 | 0.490 |
| Legume | excluded within event multiplicity | 149 | 0.232 | 0.416 |
| Legume | hog conflict | 73 | 0.158 | 0.438 |
| Legume | insufficient exact support | 133 | 0.141 | 0.353 |
| Legume | multiple hogs within species | 44 | 0.506 | 1.000 |
| Legume | primary exact unanimous | 3,012 | 0.168 | 0.336 |
| Legume | unmapped | 475 | 0.249 | 0.482 |
| Maleae (Apple/Pear) | analysed | 2,629 | 0.658 | 0.223 |
| Maleae (Apple/Pear) | not analysed | 7,884 | 0.708 | 0.290 |
| Maleae (Apple/Pear) | primary applepear one to one | 2,629 | 0.658 | 0.223 |
| Maleae (Apple/Pear) | unmapped | 7,884 | 0.708 | 0.290 |

**Table S5. Replay under alternative orthology rules.** $r$: Pearson correlation of event-wide retention percentiles; global $T$: $(n-3)$-weighted Fisher mean over reference pairs; OR: odds ratio for retention in both WGDs.

| Mapping rule | Pair | $n$ | $r$ | OR |
|---|---|---|---|---|
| primary (exact, unanimous, one-to-one) | At-$\alpha$–Grass $\rho$ | 999 | 0.187 | 2.17 |
| primary (exact, unanimous, one-to-one) | At-$\alpha$–Legume | 1,445 | 0.232 | 2.45 |
| primary (exact, unanimous, one-to-one) | Grass $\rho$–Legume | 636 | 0.195 | 2.42 |
| primary (exact, unanimous, one-to-one) | global $T$ | 3,080 | 0.210 | |
| strict ($\geq$3 species exact support) | At-$\alpha$–Grass $\rho$ | 895 | 0.197 | 2.29 |
| strict ($\geq$3 species exact support) | At-$\alpha$–Legume | 638 | 0.208 | 2.11 |
| strict ($\geq$3 species exact support) | Grass $\rho$–Legume | 286 | 0.243 | 2.81 |
| strict ($\geq$3 species exact support) | global $T$ | 1,819 | 0.208 | |
| relaxed (multiplicity averaged) | At-$\alpha$–Grass $\rho$ | 1,127 | 0.210 | 2.42 |
| relaxed (multiplicity averaged) | At-$\alpha$–Legume | 1,613 | 0.226 | 2.39 |
| relaxed (multiplicity averaged) | Grass $\rho$–Legume | 705 | 0.209 | 2.51 |
| relaxed (multiplicity averaged) | global $T$ | 3,445 | 0.217 | |
| deep one-to-one topology only | At-$\alpha$–Grass $\rho$ | 949 | 0.181 | 2.17 |
| deep one-to-one topology only | At-$\alpha$–Legume | 336 | 0.155 | 1.76 |
| deep one-to-one topology only | Grass $\rho$–Legume | 610 | 0.196 | 2.44 |
| deep one-to-one topology only | global $T$ | 1,895 | 0.182 | |
| Pentapetalae coordinate | At-$\alpha$–Legume | 1,492 | 0.230 | 2.34 |
| Pentapetalae coordinate | global $T$ | 1,492 | 0.230 | |
| primary incl. Apple/Pear | At-$\alpha$–Grass $\rho$ | 999 | 0.187 | 2.17 |
| primary incl. Apple/Pear | At-$\alpha$–Legume | 1,445 | 0.232 | 2.45 |
| primary incl. Apple/Pear | At-$\alpha$–Maleae (Apple/Pear) | 813 | 0.168 | 1.51 |
| primary incl. Apple/Pear | Grass $\rho$–Legume | 636 | 0.195 | 2.42 |
| primary incl. Apple/Pear | Grass $\rho$–Maleae (Apple/Pear) | 315 | 0.068 | 1.28 |
| primary incl. Apple/Pear | Legume–Maleae (Apple/Pear) | 532 | 0.174 | 2.06 |
| primary incl. Apple/Pear | global $T$ | 3,080 | 0.210 | |

**Table S6. Replay before and after adjustment for 76 gene properties.** Cross-fit: five-fold cross-fitted residual correlation; in-sample: ordinary least squares residual correlation with bootstrap interval; +GO ridge: cross-fitted ridge including 488 GO terms; fraction: share of between-event covariance explained in-sample after subtracting its permutation null; OR: odds ratio for retention in both WGDs, unadjusted and adjusted for covariate risk scores.

| Pair | $n$ | Unadjusted $r$ | Cross-fit | In-sample | +GO ridge | Fraction | OR | Adjusted OR |
|---|---|---|---|---|---|---|---|---|
| At-$\alpha$–Grass $\rho$ | 984 | 0.186 (0.119–0.237) | 0.122 | 0.116 (0.045–0.177) | 0.146 | 39% | 2.17 | 1.85 (1.42–2.47) |
| At-$\alpha$–Legume | 1,426 | 0.231 (0.182–0.283) | 0.156 | 0.153 (0.098–0.208) | 0.160 | 39% | 2.44 | 1.95 (1.55–2.45) |
| At-$\alpha$–Maleae (Apple/Pear) | 794 | 0.169 (0.111–0.230) | 0.114 | 0.131 (0.064–0.208) | 0.151 | 23% | 1.53 | 1.49 (1.06–2.08) |
| Grass $\rho$–Legume | 368 | 0.178 (0.081–0.275) | 0.114 | 0.082 (-0.031–0.193) | 0.138 | 48% | 2.15 | 2.09 (1.37–3.32) |
| Grass $\rho$–Maleae (Apple/Pear) | 160 | 0.078 (-0.092–0.236) | 0.045 | 0.016 (-0.228–0.364) | 0.084 | 45% | 1.54 | 1.58 (0.71–3.45) |
| Legume–Maleae (Apple/Pear) | 262 | 0.166 (0.049–0.273) | 0.115 | 0.177 (0.045–0.339) | 0.159 | -2% | 2.03 | 2.05 (1.19–3.57) |

**Table S7. Predicting retention in held-out plant WGDs.** Lineage: mean retention percentile in the other plant WGDs (no fitting). Properties: cross-fitted ridge-logistic model of 76 gene properties. Combined: properties plus lineage score. $\Delta$AUC: gain of the combined over the property model (paired bootstrap). Q4/Q1: retention in top versus bottom quartile of the lineage score.

| Target | $n$ | Prevalence | Lineage AUC | Properties AUC | Combined AUC | $\Delta$AUC | Q4/Q1 |
|---|---|---|---|---|---|---|---|
| At-$\alpha$ | 2,553 | 0.36 | 0.62 (0.60–0.65) | 0.64 (0.62–0.67) | 0.67 (0.65–0.69) | +0.027 (+0.016, +0.037) | 2.02 |
| Grass $\rho$ | 1,037 | 0.32 | 0.61 (0.58–0.65) | 0.59 (0.56–0.63) | 0.62 (0.58–0.66) | +0.026 (+0.012, +0.040) | 1.78 |
| Legume | 1,477 | 0.37 | 0.63 (0.61–0.66) | 0.69 (0.66–0.72) | 0.71 (0.68–0.73) | +0.020 (+0.008, +0.031) | 1.67 |
| Maleae (Apple/Pear) | 837 | 0.25 | 0.58 (0.54–0.63) | 0.47 (0.42–0.52) | 0.48 (0.44–0.53) | +0.011 (-0.002, +0.025) | 1.90 |

**Table S8. Cross-kingdom concordance at the Eukaryota coordinate** (TGD–Stylommatophora at the Bilateria coordinate). Retention in B given retention / loss in A; one-sided permutation $P$ for Spearman $\rho$ (10,000 permutations); latent: cross-event tetrachoric correlation disattenuated for within-event agreement (Note S5). Stylommatophora retention: both copies in at least two of three genomes.

| Pair | $n$ | $P(B \mid A)$ / $P(B \mid \bar{A})$ | Odds ratio | Spearman $\rho$ | $P$ | Latent $r$ |
|---|---|---|---|---|---|---|
| TGD–Yeast | 506 | 32% / 18% | 2.10 (1.29–3.37) | 0.15 (0.05–0.24) | 0.0010 | 0.31 (0.08–0.45) |
| At-$\alpha$–Yeast | 449 | 32% / 14% | 2.92 (1.84–4.84) | 0.23 (0.13–0.32) | $\leq$0.0001 | 0.37 (0.21–0.50) |
| Grass $\rho$–Yeast | 234 | 42% / 24% | 2.31 (1.29–4.20) | 0.21 (0.08–0.35) | 0.0008 | 0.32 (0.06–0.50) |
| Legume–Yeast | 274 | 38% / 22% | 2.21 (1.19–3.95) | 0.18 (0.05–0.31) | 0.0018 | 0.41 (0.16–0.58) |
| Maleae (Apple/Pear)–Yeast | 107 | 29% / 24% | 1.30 (0.38–3.39) | 0.01 (-0.20–0.20) | 0.4761 | -0.28 (-0.61–0.13) |
| At-$\alpha$–TGD | 348 | 24% / 21% | 1.25 (0.75–2.09) | 0.07 (-0.04–0.17) | 0.1109 | 0.13 (-0.05–0.29) |
| Grass $\rho$–TGD | 200 | 39% / 22% | 2.26 (1.17–4.44) | 0.19 (0.04–0.34) | 0.0057 | 0.28 (0.03–0.50) |
| Legume–TGD | 224 | 29% / 19% | 1.68 (0.81–3.33) | 0.14 (-0.01–0.28) | 0.0230 | 0.39 (0.10–0.62) |
| Maleae (Apple/Pear)–TGD | 116 | 21% / 26% | 0.75 (0.15–2.05) | -0.14 (-0.32–0.05) | 0.9453 | -0.29 (-0.60–0.06) |
| TGD–Stylommatophora | 146 | 65% / 52% | 1.69 (0.85–3.52) | 0.23 (0.05–0.39) | 0.0027 | |

**Table S9. Dates used for the tree in Fig. 1A.** Divergence and crown ages are TimeTree 5 medians and 95% intervals for the named species pairs (retrieved October 2026). Duplication ages are published estimates; for the apple–pear and teleost duplications the published range is shown.

| Node | Type | Species compared | Age (Ma) | Range | Source |
|---|---|---|---|---|---|
| Eukaryota (plants vs opisthokonts) | divergence | Arabidopsis thaliana vs Homo sapiens | 1,455.3 | 1,227.9–1,682.6 | TimeTree 5 (Kumar et al. 2022), median of published estimates |
| Opisthokonta (fungi vs animals) | divergence | Saccharomyces cerevisiae vs Homo sapiens | 1,217.9 | 1,113.1–1,322.6 | TimeTree 5 |
| Bilateria (deuterostomes vs protostomes) | divergence | Danio rerio vs Biomphalaria glabrata | 721.5 | 645.7–797.2 | TimeTree 5 |
| Mesangiospermae (monocots vs eudicots) | divergence | Oryza sativa vs Arabidopsis thaliana | 159.6 | 153.6–165.6 | TimeTree 5 |
| Rosids (malvids vs fabids) | divergence | Arabidopsis thaliana vs Medicago truncatula | 107.9 | 104.2–111.5 | TimeTree 5 |
| Fabids (Fabales vs Rosales) | divergence | Medicago truncatula vs Malus domestica | 97.6 | 93.3–101.9 | TimeTree 5 |
| Brassicaceae crown (sampled genomes) | crown of sampled genomes | Arabidopsis thaliana vs Aethionema arabicum | 40.6 | 37.4–43.7 | TimeTree 5 |
| Poaceae sampled crown (BOP vs PACMAD) | crown of sampled genomes | Oryza sativa vs Sorghum bicolor | 52.1 | 47.0–57.3 | TimeTree 5 |
| Papilionoideae sampled crown | crown of sampled genomes | Medicago truncatula vs Phaseolus vulgaris | 56.8 | 49.9–63.7 | TimeTree 5 |
| Maleae sampled crown | crown of sampled genomes | Malus domestica vs Pyrus communis | 15.1 | 7.3–22.9 | TimeTree 5 |
| Post-WGD Saccharomycetaceae sampled crown | crown of sampled genomes | Saccharomyces cerevisiae vs Vanderwaltozyma polyspora | 92.9 | 92.8–93.0 | TimeTree 5 |
| Teleostei sampled crown (Otocephala vs Euteleostei) | crown of sampled genomes | Danio rerio vs Tetraodon nigroviridis | 220.4 | 203.5–237.2 | TimeTree 5 |
| Stylommatophora sampled crown | crown of sampled genomes | Arion vulgaris vs Helix pomatia | 96.4 | 83.2–109.5 | TimeTree 5 |
| At-alpha WGD | duplication | Brassicaceae | 47.0 | – | Kagale et al. 2014 Plant Cell 26:2777 |
| Grass rho WGD | duplication | Poaceae | 70.0 | – | Paterson et al. 2004 PNAS 101:9903 |
| Papilionoid legume WGD | duplication | Papilionoideae | 59.0 | – | Schmutz et al. 2010 Nature 463:178 |
| Maleae (Apple/Pear) WGD | duplication | Maleae | 37.5 | 30.0–45.0 | Wu et al. 2013 Genome Res 23:396 |
| Yeast WGD | duplication | Saccharomycetaceae | 100.0 | – | Wolfe & Shields 1997 Nature 387:708 |
| Teleost genome duplication (TGD) | duplication | Teleostei | 276.7 | 267.2–286.2 | Qi et al. 2024 Genome Biol Evol 16:evae128 (earlier estimates 320-350 Ma) |
| Stylommatophora WGD | duplication | Stylommatophora | – | >96 | Shared by all sampled stylommatophorans and absent from Onchidella (Liu et al. 2021; Chen et al. 2022; McHale et al. 2025), so older than the sampled crown; a synonymous-divergence estimate of approximately 70 Ma (Liu et al. 2021) is younger than the crown and is not used |

**Table S10. Whole-genome duplications analysed.** Retained: fraction of all reconstructed ancestral loci (pillars) with both ohnologs in at least one descendant genome (apple/pear: in all five). HOGs: ancestral genes matched one-to-one at the stated level. [a]Accepted loci reconstructed in this study (618 high- and 349 moderate-confidence).

| WGD | Clade | Genomes | Pillars | Retained | HOGs matched (level) | Role |
|---|---|---|---|---|---|---|
| At-$\alpha$ | Brassicaceae | 6 | 7,243 | 0.42 | 4,934 (Mesangiospermae) | reference |
| Grass $\rho$ | Poaceae | 5 | 2,855 | 0.34 | 2,158 (Mesangiospermae) | reference |
| Legume | Papilionoideae | 4 | 3,886 | 0.37 | 2,891 (Mesangiospermae) | reference |
| Apple/Pear | Maleae (Rosaceae) | 5 | 10,513 | 0.27 | 2,629 (Mesangiospermae) | prospective target |
| Yeast | Saccharomycetaceae | 11 | 4,065 | 0.27 | 1,392 (Eukaryota) | prospective target |
| TGD | Teleostei | 8 | 5,589 | 0.41 | 1,187 (Eukaryota) | prospective target |
| Stylommatophora | land snails and slugs | 3 | 967[a] | – | 146 shared with TGD (Bilateria) | independent animal comparison |

**Table S11. Freeze records of the prespecified tests** (Note S6). For each test, the population, statistic and decision rules were fixed and the frozen artefact hashed (SHA-256) and committed before the association was computed; for the plant-to-target tests, the prediction table was committed before the target outcome was constructed (TGD, yeast) or opened (apple–pear). Hashes and commit identifiers are truncated; full values, the decision log and the frozen files are in the reproducibility package. All dates are in 2026.

| Test | Milestone | Date | Frozen before outcome | SHA-256 | Commit | Decision label |
|---|---|---|---|---|---|---|
| Reference replay and leave-one-out transfer | M4 | 28 Aug | Input hashes, seeds, statistics and gates | in freeze record | `cdaafb4ace2e`[a] | `M4_PASS_ROBUST_REPLAY_AND_TRANSFER` |
| Apple–pear | M5A–M5B | 28 Aug | Orthology map; prediction table (373 HOGs) | `632101cdf69f` | `afd9e7fbf064` | `M5B_PASS_PROSPECTIVE_EXTERNAL_VALIDATION` |
| TGD | M6B–M6C | 29 Aug | Prediction table (151 HOGs) | `9c8e0d159cf1` | `d6d30f59b3fc`[b] | `M6C_NO_SUPPORTED_CROSS_CLADE_TRANSFER` |
| Budding yeast | M6D–M6E | 29 Aug | Prediction table (186 HOGs) | `1e829dc81e09` | `b8ddd8b3867a` | `M6E_PASS_STRONG_CROSS_KINGDOM_TRANSFER` |
| TGD–snail | M11B-R | 3 Sep | Complete-case set (146 HOGs), rank hashes, gates | `b3dcbf00d66f` | in freeze record | `M11BR_POSITIVE_BUT_BELOW_STRONG_REPLAY_GATE` |
| Plant score–snail | M13B | 3 Sep | Universe (25 HOGs), statistic, gates | in freeze record | `1912845b1c89` | `M13B_POSITIVE_BUT_UNRESOLVED_CROSS_KINGDOM_REPLAY` |

[a]Commit freezing the implementation and result; input hashes were recorded before the join. [b]Made after the protocol commit `aa894827d751` and before the TGD outcome was constructed.

**Table S12. Within-genome coupling between complex partners.** Two-way fixed-effects (gene, genome) linear probability model of a gene's two-copy state on the fraction of its complex partners kept in the same genome (partners within 20 genes in the ancestral order and paralogous lineages of one eukaryotic gene excluded). Null A: partner sets permuted among complex-member genes (2,000 permutations); null B: members replaced by random genes of the same WGD with the same occupancy (500). Informative: genes with both states among the genomes. 95% CI: 2,000 (yeast) or 1,000 bootstrap resamples of focal genes. Pooled: Stouffer combination of $z_A$ over the five non-yeast WGDs.

| WGD | Complexes | Genomes | Genes | Informative | Pairs | $\theta$ | 95% CI | Null A | $z_A$ | $P_A$ | Null B | $z_B$ |
|---|---|---|---|---|---|---|---|---|---|---|---|---|
| Yeast | curated | 11 | 697 | 188 | 5,881 | 0.31 | 0.22 to 0.41 | $-0.005 \pm 0.040$ | 8.1 | 0.0005 | $0.124 \pm 0.052$ | 3.7 |
| TGD | curated | 8 | 513 | 135 | 2,377 | –0.06 | –0.15 to 0.03 | $-0.005 \pm 0.041$ | –1.3 | 0.909 | $-0.020 \pm 0.050$ | –0.7 |
| At-$\alpha$ | curated | 6 | 846 | 339 | 46,386 | –0.07 | –0.15 to 0.01 | $-0.007 \pm 0.048$ | –1.3 | 0.894 | $-0.014 \pm 0.046$ | –1.1 |
| At-$\alpha$ | GO component | 6 | 169 | 79 | 2,311 | 0.04 | – | $-0.205 \pm 0.302$ | 0.8 | 0.223 | $-0.140 \pm 0.281$ | 0.6 |
| Grass $\rho$ | curated | 5 | 342 | 139 | 9,183 | –0.04 | –0.21 to 0.10 | $-0.012 \pm 0.068$ | –0.4 | 0.680 | $-0.015 \pm 0.063$ | –0.4 |
| Grass $\rho$ | GO component | 5 | 38 | 15 | 120 | –0.58 | – | $-0.131 \pm 0.247$ | –1.8 | 0.952 | $-0.177 \pm 0.314$ | –1.3 |
| Legume | curated | 4 | 385 | 146 | 11,783 | –0.05 | –0.12 to 0.01 | $-0.011 \pm 0.082$ | –0.4 | 0.686 | $-0.011 \pm 0.069$ | –0.5 |
| Legume | GO component | 4 | 50 | 15 | 261 | –0.20 | – | $-0.190 \pm 0.311$ | –0.0 | 0.546 | $-0.293 \pm 0.413$ | 0.2 |
| Maleae (Apple/Pear) | curated | 5 | 194 | 201 | 3,734 | –0.12 | –0.29 to 0.04 | $-0.026 \pm 0.096$ | –1.0 | 0.837 | $-0.042 \pm 0.087$ | –0.9 |
| Maleae (Apple/Pear) | GO component | 5 | 21 | 16 | 69 | 0.25 | – | $-0.249 \pm 0.350$ | 1.4 | 0.071 | $-0.399 \pm 0.380$ | 1.7 |
| Non-yeast WGDs pooled | curated | | | | | | | | –2.0 | | | |

**Table S13. Yeast coupling: robustness, specificity, complex classes and power.** $\theta$ and $z$ as in Table S12; neighbour window $W$ in genes of the ancestral order; class rows restrict partners to the named class with a null of random member sets of the same size. Proteasome and chaperonin rows are omitted because almost no genes vary in state across genomes.

| Analysis | $\theta$ | Genes | $z$ | $P$ |
|---|---|---|---|---|
| all curated, W=0 | 0.32 | 699 | 8.2 | 0.001 |
| all curated, W=20 | 0.31 | 697 | 8.4 | 0.001 |
| all curated, W=50 | 0.32 | 695 | 8.2 | 0.001 |
| all curated, W=100 | 0.31 | 693 | 7.8 | 0.001 |
| excluding ribosomal proteins | 0.09 | 573 | 2.5 | 0.008 |
| ribosomal proteins only | 0.77 | 123 | 3.8 | 0.001 |
| source CYC2008 only | 0.43 | 479 | 9.1 | 0.001 |
| source CORUM only | 0.23 | 499 | 5.0 | 0.001 |
| source Complex Portal only | 0.36 | 600 | 9.0 | 0.001 |
| drop genome Candida glabrata | 0.27 | 697 | 6.5 | 0.0025 |
| drop genome Kazachstania africana | 0.33 | 697 | 7.7 | 0.0025 |
| drop genome Kazachstania naganishii | 0.33 | 697 | 8.7 | 0.0025 |
| drop genome Naumovozyma castellii | 0.31 | 697 | 8.5 | 0.0025 |
| drop genome Naumovozyma dairenensis | 0.33 | 697 | 8.8 | 0.0025 |
| drop genome Saccharomyces cerevisiae | 0.32 | 697 | 8.7 | 0.0025 |
| drop genome Saccharomyces kudriavzevii | 0.32 | 697 | 8.6 | 0.0025 |
| drop genome Saccharomyces uvarum | 0.32 | 697 | 9.4 | 0.0025 |
| drop genome Tetrapisispora blattae | 0.27 | 697 | 6.5 | 0.0025 |
| drop genome Tetrapisispora phaffii | 0.32 | 697 | 8.6 | 0.0025 |
| drop genome Vanderwaltozyma polyspora | 0.33 | 697 | 8.3 | 0.0025 |
| relation: physical complex partners (no ribosome) | 0.09 | 568 | 2.5 | 0.008 |
| relation: same GO process, not in a shared complex | −0.09 | 350 | −0.8 | 0.786 |
| relation: both complex and process | 0.22 | 89 | 3.1 | 0.007 |
| class: cytosolic ribosome | 0.78 | 127 | 3.5 | 0.001 |
| class: RNA polymerase | −0.06 | 26 | 0.8 | 0.179 |
| class: translation initiation | 0.52 | 41 | 2.6 | 0.004 |
| class: spliceosome / snRNP | 0.95 | 39 | 3.5 | 0.001 |
| class: mediator / transcription | 0.08 | 31 | 1.1 | 0.094 |
| class: mitochondrial respiratory | −0.21 | 20 | 0.3 | 0.394 |
| leave out class: cytosolic ribosome | 0.09 | 568 | 2.3 | 0.014 |
| leave out class: spliceosome / snRNP | 0.31 | 654 | 8.3 | 0.002 |
| leave out class: translation initiation | 0.31 | 651 | 8.0 | 0.002 |
| leave out class: proteasome | 0.31 | 663 | 8.9 | 0.002 |
| leave out class: RNA polymerase | 0.31 | 670 | 8.0 | 0.002 |
| leave out class: mediator / transcription | 0.32 | 668 | 8.5 | 0.002 |
| leave out class: mitochondrial respiratory | 0.33 | 677 | 8.7 | 0.002 |
| bootstrap 95% CI, all complexes: 0.22 to 0.41 | 0.31 | – | – | – |
| bootstrap 95% CI, ribosome excluded: −0.01 to 0.19 | 0.09 | – | – | – |
| sister pairs, recent co-loss: partners lost in the same sister 500 vs other sister 238 (68% vs null 53 $\pm$ 7%) | – | – | 2.1 | 0.027 |
| sister pairs, precedence: earlier partner loss and later gene loss (300 gene-pair observations) | 0.21 | – | 0.7 | 0.221 |
| dose moderation, MoBY-CEN extra-copy cost: high 0.55 vs low 0.29 (difference, one-sided permutation) | 0.26 | 260 | – | 0.087 |
| dose moderation, gTOW copy-number-limit sensitivity: high 0.35 vs low 0.28 (difference, one-sided permutation) | 0.07 | 812 | – | 0.223 |
| power: random subsets of 4 genomes (40); detected at $P \leq 0.05$ in 92% | 0.34 | – | 6.3 | – |
| power: random subsets of 5 genomes (40); detected at $P \leq 0.05$ in 95% | 0.32 | – | 6.7 | – |
| power: random subsets of 6 genomes (40); detected at $P \leq 0.05$ in 100% | 0.32 | – | 6.8 | – |

**Table S14. Within-genome coupling between neighbouring genes.** Two-way fixed-effects model of a gene's two-copy state on the mean state of the genes at the stated distance (both sides) in the ancestral pillar order, using all POInT pillars; null: genes placed in random order (100 permutations).

| WGD | Analysis | Pillars | $\theta$ | $z$ |
|---|---|---|---|---|
| Maleae (Apple/Pear) | distance 1 | 10,513 | 0.315 | 38.2 |
| Maleae (Apple/Pear) | distance 2 | 10,513 | 0.240 | 34.4 |
| Maleae (Apple/Pear) | distance 5 | 10,513 | 0.179 | 24.9 |
| Maleae (Apple/Pear) | distance 20 | 10,513 | 0.090 | 14.2 |
| Maleae (Apple/Pear) | distance 100 | 10,513 | 0.025 | 3.5 |
| Maleae (Apple/Pear) | distance 1, drop Malus baccata | 10,513 | 0.356 | 34.1 |
| Maleae (Apple/Pear) | distance 1, drop Malus domestica | 10,513 | 0.334 | 37.5 |
| Maleae (Apple/Pear) | distance 1, drop Malus sieversii | 10,513 | 0.330 | 38.1 |
| Maleae (Apple/Pear) | distance 1, drop Pyrus bretschneideri | 10,513 | 0.174 | 21.1 |
| Maleae (Apple/Pear) | distance 1, drop Pyrus communis | 10,513 | 0.359 | 42.4 |
| At-$\alpha$ | distance 1 | 7,243 | 0.076 | 9.4 |
| At-$\alpha$ | distance 2 | 7,243 | 0.011 | 1.6 |
| At-$\alpha$ | distance 5 | 7,243 | 0.017 | 2.0 |
| At-$\alpha$ | distance 20 | 7,243 | −0.006 | −0.8 |
| At-$\alpha$ | distance 100 | 7,243 | 0.006 | 0.6 |
| At-$\alpha$ | distance 1, drop Aethionema arabicum | 7,243 | 0.055 | 6.1 |
| At-$\alpha$ | distance 1, drop Arabidopsis lyrata | 7,243 | 0.061 | 7.0 |
| At-$\alpha$ | distance 1, drop Arabidopsis thaliana | 7,243 | 0.079 | 10.4 |
| At-$\alpha$ | distance 1, drop Capsella rubella | 7,243 | 0.085 | 10.9 |
| At-$\alpha$ | distance 1, drop Eutrema salsugineum | 7,243 | 0.084 | 8.6 |
| At-$\alpha$ | distance 1, drop Schrenkiella parvula | 7,243 | 0.088 | 9.7 |
| Grass $\rho$ | distance 1 | 2,855 | 0.101 | 7.5 |
| Grass $\rho$ | distance 2 | 2,855 | 0.042 | 3.3 |
| Grass $\rho$ | distance 5 | 2,855 | 0.015 | 0.9 |
| Grass $\rho$ | distance 20 | 2,855 | −0.002 | −0.1 |
| Grass $\rho$ | distance 100 | 2,855 | −0.004 | −0.2 |
| Grass $\rho$ | distance 1, drop Brachypodium distachyon | 2,855 | 0.071 | 4.7 |
| Grass $\rho$ | distance 1, drop Oropetium thomaeum | 2,855 | 0.123 | 9.2 |
| Grass $\rho$ | distance 1, drop Oryza sativa | 2,855 | 0.111 | 7.2 |
| Grass $\rho$ | distance 1, drop Setaria italica | 2,855 | 0.088 | 6.4 |
| Grass $\rho$ | distance 1, drop Sorghum bicolor | 2,855 | 0.113 | 8.0 |
| Legume | distance 1 | 3,886 | 0.167 | 13.5 |
| Legume | distance 2 | 3,886 | 0.049 | 3.7 |
| Legume | distance 5 | 3,886 | 0.007 | 0.6 |
| Legume | distance 20 | 3,886 | −0.044 | −3.4 |
| Legume | distance 100 | 3,886 | 0.014 | 1.0 |
| Legume | distance 1, drop Cajanus cajan | 3,886 | 0.168 | 12.0 |
| Legume | distance 1, drop Cicer arietinum | 3,886 | 0.142 | 9.3 |
| Legume | distance 1, drop Medicago truncatula | 3,886 | 0.184 | 11.7 |
| Legume | distance 1, drop Phaseolus vulgaris | 3,886 | 0.173 | 10.0 |
| Yeast | distance 1 | 4,065 | 0.022 | 2.3 |
| Yeast | distance 2 | 4,065 | 0.002 | 0.2 |
| Yeast | distance 5 | 4,065 | 0.005 | 0.4 |
| Yeast | distance 20 | 4,065 | 0.000 | 0.1 |
| TGD | distance 1 | 5,589 | 0.154 | 19.1 |
| TGD | distance 2 | 5,589 | 0.089 | 11.7 |
| TGD | distance 5 | 5,589 | 0.044 | 6.4 |
| TGD | distance 20 | 5,589 | 0.016 | 2.0 |

**Table S15. What carries the shared propensity.** Spearman $\rho$ of gene features with the posterior shared ($L$) and WGD-specific ($D$) propensities (plant genes in at least two WGDs); associations of pre-duplication sequence constraint and length (outgroup alignments) and laboratory dose measures (gTOW copy-number-limit sensitivity, MoBY-CEN extra-copy cost) with retention after the yeast WGD and TGD; and partial Spearman correlations (rank residuals, 1,000 bootstrap resamples) of the plant shared propensity with yeast or TGD retention.

| Analysis | Feature or conditioning | $n$ | Estimate | Comparison |
|---|---|---|---|---|
| shared vs WGD-specific propensity (plants) | atlas expr magnitude | 316 | 0.17 | 0.04 |
| shared vs WGD-specific propensity (plants) | publications log1p | 2,621 | 0.15 | 0.07 |
| shared vs WGD-specific propensity (plants) | dosage cis deviation | 2,552 | 0.13 | 0.06 |
| shared vs WGD-specific propensity (plants) | duplication depth | 3,271 | 0.12 | 0.05 |
| shared vs WGD-specific propensity (plants) | n go experimental | 2,621 | 0.11 | 0.04 |
| shared vs WGD-specific propensity (plants) | protein length aln | 309 | −0.10 | 0.01 |
| shared vs WGD-specific propensity (plants) | ppi degree | 2,621 | 0.10 | 0.05 |
| shared vs WGD-specific propensity (plants) | atlas expr breadth | 307 | 0.10 | −0.01 |
| shared vs WGD-specific propensity (plants) | dosage trans response | 2,542 | 0.09 | 0.03 |
| shared vs WGD-specific propensity (plants) | complex conserved | 1,493 | 0.09 | 0.04 |
| shared vs WGD-specific propensity (plants) | seq constraint | 309 | 0.09 | 0.02 |
| shared vs WGD-specific propensity (plants) | phylostratum | 3,271 | 0.06 | 0.02 |
| shared vs WGD-specific propensity (plants) | dosage tetraploid deviation | 2,553 | −0.05 | −0.03 |
| shared vs WGD-specific propensity (plants) | family n angiosperm lineages | 3,271 | 0.05 | 0.02 |
| shared vs WGD-specific propensity (plants) | essential | 2,621 | −0.04 | −0.01 |
| shared vs WGD-specific propensity (plants) | protein abundance log10ppm | 1,909 | −0.01 | −0.01 |
| shared vs WGD-specific propensity (plants) | any mutant phenotype | 2,621 | 0.01 | 0.01 |
| shared vs WGD-specific propensity (plants) | expr seedling log2rpkm | 2,616 | −0.01 | −0.00 |
| retention after the Yeast WGD | constraint | 1,163 | 0.26 ($P < 0.0001$) | 0.17 (no ribosomal proteins) |
| retention after the Yeast WGD | log length | 1,163 | −0.10 ($P = 0.0004$) | 0.00 (no ribosomal proteins) |
| retention after the Yeast WGD | complex | 1,392 | 0.03 ($P = 0.310$) | −0.04 (no ribosomal proteins) |
| retention after the Yeast WGD | ribosomal | 1,392 | 0.24 ($P < 0.0001$) | – (no ribosomal proteins) |
| retention after the Yeast WGD | extra copy cost | 450 | 0.08 ($P = 0.112$) | −0.03 (no ribosomal proteins) |
| retention after the Yeast WGD | copy limit sensitivity | 1,380 | 0.16 ($P < 0.0001$) | 0.14 (no ribosomal proteins) |
| retention after the TGD WGD | constraint | 615 | 0.17 ($P < 0.0001$) | 0.17 (no ribosomal proteins) |
| retention after the TGD WGD | log length | 615 | −0.04 ($P = 0.370$) | −0.02 (no ribosomal proteins) |
| retention after the TGD WGD | complex | 1,187 | −0.05 ($P = 0.119$) | −0.03 (no ribosomal proteins) |
| retention after the TGD WGD | ribosomal | 1,187 | −0.07 ($P = 0.012$) | – (no ribosomal proteins) |
| retention after the TGD WGD | extra copy cost | 180 | 0.11 ($P = 0.158$) | 0.16 (no ribosomal proteins) |
| retention after the TGD WGD | copy limit sensitivity | 561 | 0.04 ($P = 0.407$) | 0.05 (no ribosomal proteins) |
| plant shared propensity vs Yeast retention | conditioned on none | 630 | 0.205 (0.12–0.28) | |
| plant shared propensity vs Yeast retention | conditioned on constraint | 630 | 0.143 (0.06–0.23) | |
| plant shared propensity vs Yeast retention | conditioned on length | 630 | 0.192 (0.11–0.27) | |
| plant shared propensity vs Yeast retention | conditioned on ribosomal | 630 | 0.190 (0.11–0.26) | |
| plant shared propensity vs Yeast retention | conditioned on copy-limit sensitivity | 630 | 0.202 (0.12–0.28) | |
| plant shared propensity vs Yeast retention | conditioned on constraint+length+ribosomal | 630 | 0.137 (0.06–0.22) | |
| plant shared propensity vs Yeast retention | conditioned on all four | 630 | 0.137 (0.05–0.22) | |
| plant shared propensity vs Yeast retention | conditioned on (constraint vs plant shared) | 630 | 0.249 (0.18–0.32) | |
| plant shared propensity vs TGD retention | conditioned on none | 162 | 0.078 (−0.09–0.23) | |
| plant shared propensity vs TGD retention | conditioned on constraint | 162 | 0.040 (−0.15–0.21) | |
| plant shared propensity vs TGD retention | conditioned on length | 162 | 0.062 (−0.09–0.21) | |
| plant shared propensity vs TGD retention | conditioned on ribosomal | 162 | 0.108 (−0.04–0.27) | |
| plant shared propensity vs TGD retention | conditioned on copy-limit sensitivity | 162 | 0.077 (−0.08–0.24) | |
| plant shared propensity vs TGD retention | conditioned on constraint+length+ribosomal | 162 | 0.056 (−0.12–0.23) | |
| plant shared propensity vs TGD retention | conditioned on all four | 162 | 0.056 (−0.12–0.22) | |
| plant shared propensity vs TGD retention | conditioned on (constraint vs plant shared) | 162 | 0.276 (0.13–0.41) | |
